# Prothrombinase processivity is conferred by substrate allostery

**DOI:** 10.64898/2026.01.12.699050

**Authors:** Fatma Işık Üstok, Alexandre Faille, Alan J. Warren, James A. Huntington

**Author notes:** Address correspondence to James A. Huntington, Cambridge Institute for Medical Research, Department of Haematology, University of Cambridge, The Keith Peters Building, Hills Road, Cambridge, CB2 0XY, United Kingdom. Authors contributed equally (Fatma Işık Üstok, Alexandre Faille). Data Sharing: Emails to corresponding author.

## Abstract

Conversion of prothrombin to thrombin occurs in the final step of the blood coagulation cascade and depends on association of the serine protease, factor (f) Xa, and the cofactor fVa on activated cell surfaces to form the prothrombinase complex. Prothrombinase cleaves prothrombin at two sites in a processive manner ∼500,000-times faster than fXa on its own. How fVa confers rapid and processive cleavage of prothrombin is an enzymatical mystery with profound consequence. We created a variant of fXa that binds to fVa with high affinity in the absence of phospholipids that preserves the activity of wild-type prothrombinase, and recently reported on the cryo-EM structure of the complex. It revealed an extensive interface between the two proteins, including a critical interaction between the first acidic region C-terminal to the A2 domain of fVa (the N-terminal portion of the a2-loop) with the heparin binding site of fXa. Here we present the cryo-EM structures of prothrombinase bound to prothrombin and the intermediate meizothrombin, both to 3.1 Å resolution. The prothrombin complex revealed a surprising interaction between the second acidic region of the a2-loop with exosite I of prothrombin, accounting for 70% of the total buried surface area. Cleavage at Arg320 triggers the zymogen-to-protease conformational change in meizothrombin which alters all domain-domain and fVa interactions, and results in the presentation of the second cleavage site (Arg271) for processing. Together, these structures reveal a remarkable enzymatic mechanism that depends on the active participation of the substrate itself, and introduce the new paradigm of substrate allostery.

## INTRODUCTION

The serine protease thrombin is the enzyme that clots blood; it is the only endogenous factor capable of cleaving fibrinogen into fibrin and is the most potent activator of platelets^1^. Inappropriate or excessive thrombin generation is the cause of all forms of thrombosis, and insufficient thrombin generation results in bleeding^2^. Its zymogen precursor prothrombin is comprised of a gamma carboxyglutamic acid (Gla) domain, two kringle (K) domains and a serine protease (SP) domain, with the Gla-K1 domains and the K2-SP domains each forming rigid units separated by a flexible 26-residue linker^3^ (Supp. Fig. S1A). Prothrombin circulates predominately in a closed configuration with the Gla-K1 unit (also known as fragment 1 or F1) interacting with the SP domain^4,5^. Conversion of prothrombin to thrombin requires proteolytic cleavage at two sites, after Arg271 and Arg320 (Supp. Fig. S1B), by the enzyme complex prothrombinase, comprised of a serine protease, factor (f) Xa, and a large cofactor, fVa^6,7^. To limit thrombin production to sites of vascular damage, prothrombinase can only assemble on activated phospholipid (PL) surfaces that are rich in phosphatidyl serine^8,9^. Factor Xa on its own can cleave prothrombin at both sites, but at rates too slow to be of physiological relevance and in an order (Arg271 first) that separates the membrane-binding Gla domain and two K domains (F1.2) from the zymogen form of the SP domain (Prethrombin-2 or Pre-2). Assembly of prothrombinase accelerates cleavage of prothrombin by ∼500,000-fold and enforces initial cleavage at Arg320, producing the active, membrane-anchored form meizothrombin as the intermediate. Each prothrombinase complex can convert over 100 molecules of prothrombin to thrombin per second in a processive manner, without apparent dissociation and reassociation of the intermediate^7^. How efficiency and processivity are conferred by the cofactor fVa lies at the heart of the process of blood coagulation and remains largely unresolved.

Factor Xa is comprised of an N-terminal Gla domain, two epidermal growth factor-like (EGF) domains and an SP domain^10^. The Gla and EGF1 domains associate into an intimate unit connected to the EGF2 domain by a 7-residue linker. The EGF2 domain forms extensive non-covalent contacts with the SP domain and are also connected by a disulfide bond. Factor Xa is fully active against peptidyl substrates and is not allosterically activated by fVa binding. Factor V circulates as a procofactor comprised of three A domains (∼300 residues each), an unstructured B-domain of 889 residues and two small membrane-binding C domains (155 residues each)^11^. The presence of the B-domain renders the pro-cofactor unable to bind to fXa. Activation to the two-chain fVa is achieved through excision of the B-domain by thrombin or fXa cleavage after Arg709 and Arg1545 (A1-A2 is the heavy chain and A3-C1-C2 is the light chain).

We previously created a variant of fXa with 17 mutations (M17) to the EGF2 and SP domains that conferred high affinity for fVa in the absence of PL, and which together recapitulated the prothrombin processing activity of fully-assembled wild-type prothrombinase^12^. We then used the M17-fVa complex to determine the structure of prothrombinase by cryogenic electron microscopy (cryo-EM)^13^. It revealed an extensive interface involving the A2 and A3 domains of fVa and the SP and EGF2 domains of fXa, driven in large part through electrostatic complementarity. One-third of the interface was contributed by the unstructured region that protrudes from the C-terminus of the A2 domain to the cleavage-activation site at Arg709 (657-709; Supp. Fig. S1C), referred to as the a2-loop. The a2-loop contains two highly acidic regions that contain sulphated tyrosines: the N-terminal region 659DDDEDSY*EIFE669 and the C-terminal region 686EPEDEESDADY*DY* (* denotes sulphation). The N-terminal region of the a2-loop was found to interact with the heparin binding site of fXa and the C-terminal region was not defined in the map of the prothrombinase complex.

Although the structure of the M17-prothrombinase complex advanced our understanding of prothrombinase assembly, it did little to address the fundamental questions regarding the cofactor function of fVa, namely: how is initial cleavage at Arg320 enforced; how is Arg271 presented to the active site of fXa after cleavage of Arg320; what conformational rearrangements ensure processivity (i.e. presentation of second cleavage site without dissociation of the intermediate); and how is high catalytic efficiency achieved? To address these issues we solved the cryo-EM structures of M17-prothrombinase with the substrate prothrombin and with the intermediate meizothrombin to 3.1 Å resolution. The structures reveal a multi-step mechanism driven by allosteric conformational rearrangements to the substrate and its interaction with prothrombinase, providing a surprising and satisfying answer to the questions regarding the cofactor function of fVa in thrombin generation. In this mechanism, the substrate is not a passive object acted upon by the enzyme, but actively participates in its own processing.

## RESULTS

### Cryo-EM structure of the prothrombinase-prothrombin complex

The cryo-EM map exhibited features consistent with an overall resolution of 3.1 Å, with excellent coverage of all domains for all three proteins, signal corresponding to glycosylation, and most side chains resolved (Fig. 1). The prothrombinase component was essentially identical to our previous structure of the apo-enzyme complex^13^, with a root-mean square deviation (RMSD) of 0.64 Å for 1452 equivalent Cα atoms and a conserved fVa-fXa interface (Supp. Tables S4 and S5) that buries a total of 4,973 Å^2^. Prothrombin was found exclusively in the ‘closed’ conformation^4,5^, with the F1 fragment interacting with the active site of the SP domain, and no evidence in any of the classes of the open form where the linker between the K1 and K2 domains is extended. Nevertheless, the Gla-domain of prothrombin is roughly co-planar with the membrane-binding domains of prothrombinase, enabling PL interaction for the ternary complex. The complex between prothrombin and prothrombinase is ‘productive’, with the 320-loop bound in the active site of fXa as a normal substrate (Supp. Fig. S5), and therefore represents the recognition complex of the initial cleavage event (note that the catalytic serine of fXa was mutated to alanine to prevent hydrolysis). The interface between prothrombin and fVa buries a total of 1,780 Å^2^ and involves two principal regions: the stretch between the A1 and A2 domains of fVa (313-320; the a1-loop) with the loop preceding Arg271 of prothrombin (253-266; the 271-loop); and the C-terminal region of the a2-loop of fVa with exosite I of prothrombin (Supp. Table S6). Surprisingly, the a2-loop accounts for 71% of the total buried surface area (BSA) between fVa and prothrombin, underscoring the importance of this contact for substrate recognition. Similarly, the fXa-prothrombin interface (total BSA of 1,458 Å^2^) is dominated by the interaction of the substrate loop (317IEGR-IV322) with the active site of fXa (Supp. Tables S7 and S8), accounting for 77% of the total BSA. The docking of prothrombin to prothrombinase therefore involves loops interacting with loops or loops interacting with an ordered domain. There are no substantive interactions involving ordered domains of prothrombin with ordered domains of prothrombinase. This ‘light-touch’ interaction is typical of a substrate, where stable complex formation would inhibit turnover. Also consistent with an efficient substrate-enzyme interaction is an interface dominated by electrostatics (Fig. 2A and 2B). This is exemplified by the basic a1-loop of fVa (303KKTRNLKKITREQRRHMKR321) interacting with the acidic 271-loop of prothrombin (249EEAVEEETGDGLDEDSDRAIE269) (Fig. 2C) and the acidic a2-loop of fVa (685-709) interacting with the basic exosite I of prothrombin (Fig. 2D).

**Figure 1.**
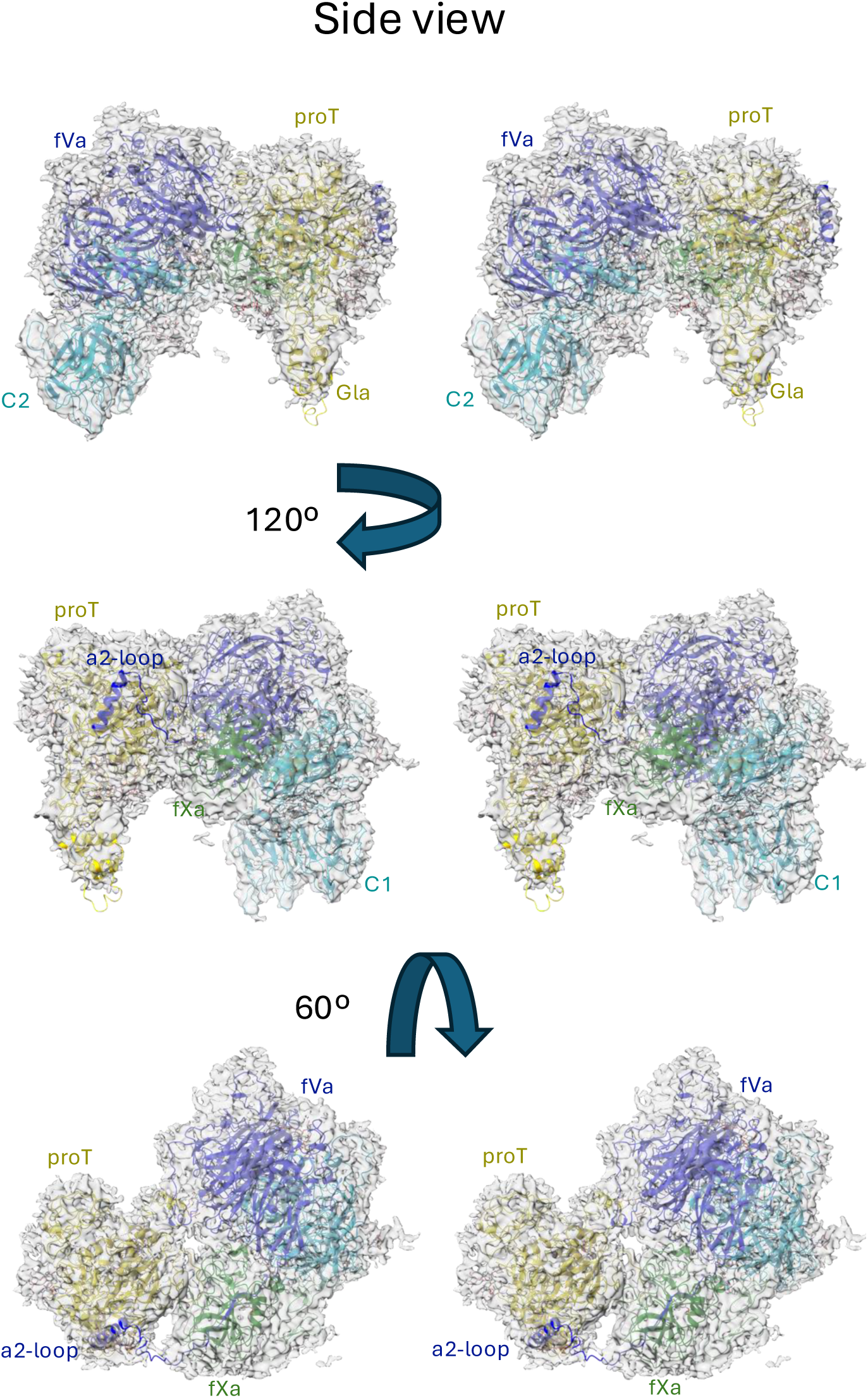
Stereo views of the structure of the prothrombinase-prothrombin complex with cryo-EM map. Three views of the final coordinates of fVa, fXa and prothrombin with surounding map are shown. The heavy chain of fVa (A1-A2 domains, including the a2-loop) is colored blue; the light chain of fVa (A3-C1-C2 domains) is colored light blue; the EGF2 and SP domains of the fXa are colored light green and green, respectively; prothrombin is colored yellow; glycosylation is indicated by grey sticks. Molecules and domains are labeled for clarity, and the figure was made using ChimeraX.

**Figure 2.**
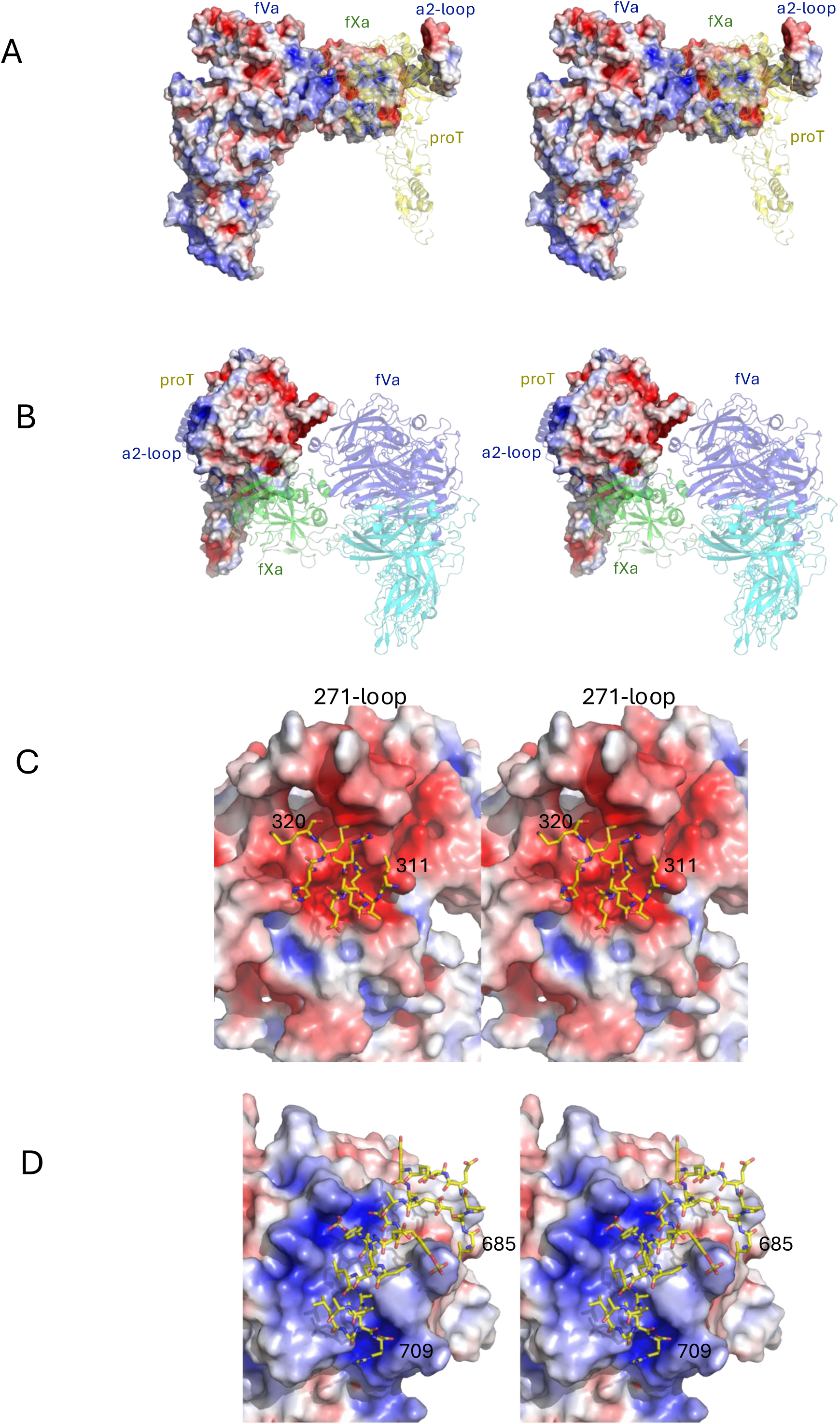
Stereo views of electrostatic surfaces of prothrombinase and prothrombin. (A) The surface of prothrombinase is colored blue for positive and red for negative electrostatic potential. Prothrombin is depicted as a semitransparent yellow cartoon. A large basic surface is presented to prothrombin by the body of fVa, and an acidic surface is presented with the a2-loop. (B) The complex is reoriented, with prothrombinase shown as semitransparent cartoons colored as in Figure 1. The electrostatic surface of prothrobin is complementary to the surfaces presented by prothrombinase, with a large acidic patch facing the body of fVa and the basic exosite I of prothrombin interacting with the acidic a2-loop C-terminus. Domains are labeled for clarity. (C) Close up of the electrostatic surface of prothrombin with the basic a1-loop of fVa (the loop connecting the A1 and A2 domains) depicted as yellow sticks. (D) Close up of the electrostatic surface of prothrombin with the a2-loop depicted as yellow sticks.

### The a2-loop:prothrombin interaction

The interaction between the C-terminal region of the a2-loop and prothrombin was unanticipated, and even more surprising was its contribution of more than 70% to the total interaction interface between fVa and prothrombin. Although this region has been implicated in prothrombin processing, the assumption was that it interacted with fXa^14,15^. We were able to build the a2-loop onto prothrombin using the cryo-EM map on its own, however, due to the unexpected nature of this interaction and to test the accuracy of its placement, we assessed the interaction computationally and structurally. AlphaFold^16^ was run using the sequences of prethrombin-2 (Pre-2), the zymogen form of the SP domain of prothrombin, and a C-terminal a2-loop peptide. The 5 outputs were identical from 690-709, with a prediction of high confidence (pIDTT>70; Supp. Fig. S6 and S7A). We also obtained a 2.8 Å crystal structure of the peptide with Pre-2 (Supp. Fig. S7B). The AlphaFold result and the crystal structure correspond well to the cryo-EM structure (Supp. Fig. S7C).

### Cryo-EM structure of the prothrombinase-meizothrombin complex

We obtained a map of the prothrombinase-meizothrombin complex to 3.1 Å resolution, employing the same conditions used for the prothrombin complex. Two surprises were immediately evident—the F1 fragment of meizothrombin and the C-terminal region of the a2-loop of fVa were both absent from the map. No particle subset possessed either feature, nor did they appear as weak signal captured using a low-resolution filter. Otherwise, the map covers all domains and is of similar quality and resolution as the map with prothrombin (Fig. 3). Importantly, the complex is productive, with the map providing clear evidence that the 271-loop is presented as a substrate in the active site of fXa (Supp. Fig. S8A), in a manner similar to the 320-loop of prothrombin (Supp. Fig. S8B). The prothrombinase component of the meizothrombin complex is again identical to apo prothrombinase (RMSD of 0.8 Å for 1455 Cα atoms) and, other than loss of the C-terminus of the a2-loop, is also identical to the structure with prothrombin (RMSD 0.61 Å). However, the two domains observable for meizothrombin (K2 and SP) have shifted substantially and make entirely new contacts with prothrombinase. The contact between meizothrombin and fVa is limited in nature with a total BSA of 327 Å^2^, involving only the 271-loop of meizothrombin (contacting residues 261-266) with the a1-loop of fVa (contacting residues 314-318; Supp. Table S9). The contact between meizothrombin and fXa is more extensive, with a total BSA of 2,187 Å^2^, half of which involves the substrate loop (268IEGR-TA273) with the active site of fXa (Supp. Table S10). The other interactions between meizothrombin and fXa can be considered exosite interactions, and primarily consist of the remnant of the 320-loop of meizothrombin (310-320) with the 150-loop of fXa (145-150). The only contacts involving the SP domain of meizothrombin also engage the 150-loop of fXa (Supp. Table S11). Despite the large shift in position of meizothrombin relative to that of prothrombin, the interface still exhibits a highly complementary electrostatic nature (Fig. 4).

**Figure 3.**
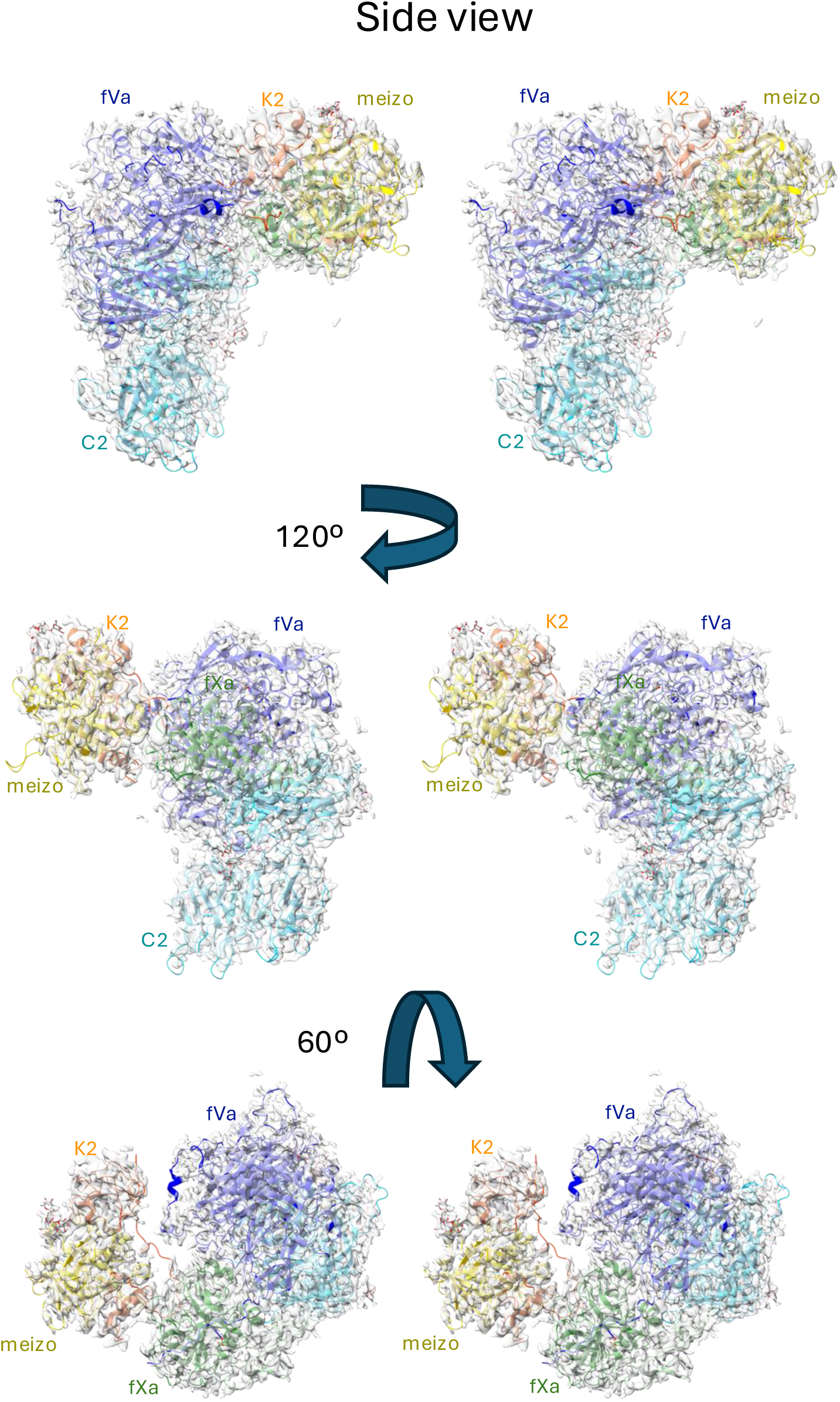
Stereo views of the structure of the prothrombinase-meizothrombin complex with cryo-EM map. Three views of the final coordinates of fVa, fXa and meizothrombin with surounding map are shown, colored and oriented roughly as in Figure 1. The light chain of meizothrombin, including the K2 domain, is colored orange to distinguish it from the SP domain which is in yellow.

**Figure 4.**
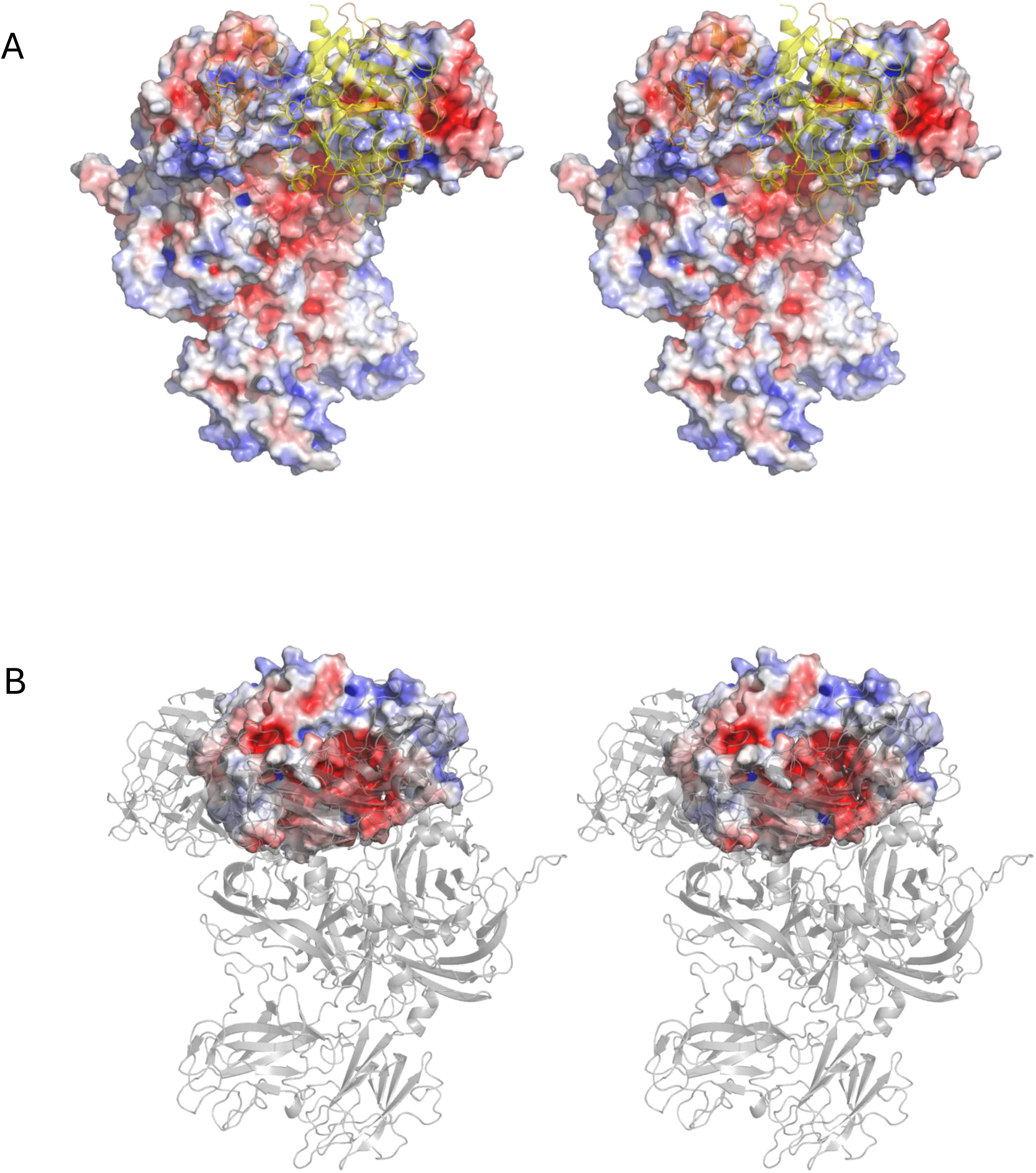
Stereo views of electrostatic surfaces of prothrombinase and meizothrombin. (A) The surface of prothrombinase is colored blue for positive and red for negative electrostatic potential. Meizothrombin is depicted as a semitransparent cartoon, colored yellow for the SP domain and orange for the light chain, including the K2 domain. The interaction surface on prothrombinase is overwhelmingly basic. (B) The complex is reoriented to look through the semitransparent cartoon representation of prothrombinase (gray) at the acidic electrostatic surface of meizothrombin.

### The prothrombin to meizothrombin conformational change

It is worth reiterating that the prothrombinase complex remains unaltered during prothrombin binding and processing (Supp. Fig. S9), and that only the substrate undergoes conformational rearrangements. Focusing on the SP domain, conversion of prothrombin to meizothrombin is associated with a 50° rotation and a 10 Å shift (Fig. 5 and Supp. Movies S1 and S2). This reorientation is triggered by the zymogen-to-protease conformational change^17^ in the SP domain following insertion of the new N-terminus (321IVEG324; 16-19 in chymotrypsin numbering) into the core of the SP domain. Activation of meizothrombin results in dramatic alterations to the shape and properties of the active site region (Fig. 6A) and converts extensive interactions with K1(Supp. Table S12) into severe clashes (Supp. Table S13), explaining the loss of contact and conversion to the open configuration, consistent with recent findings^18^.

**Figure 5.**
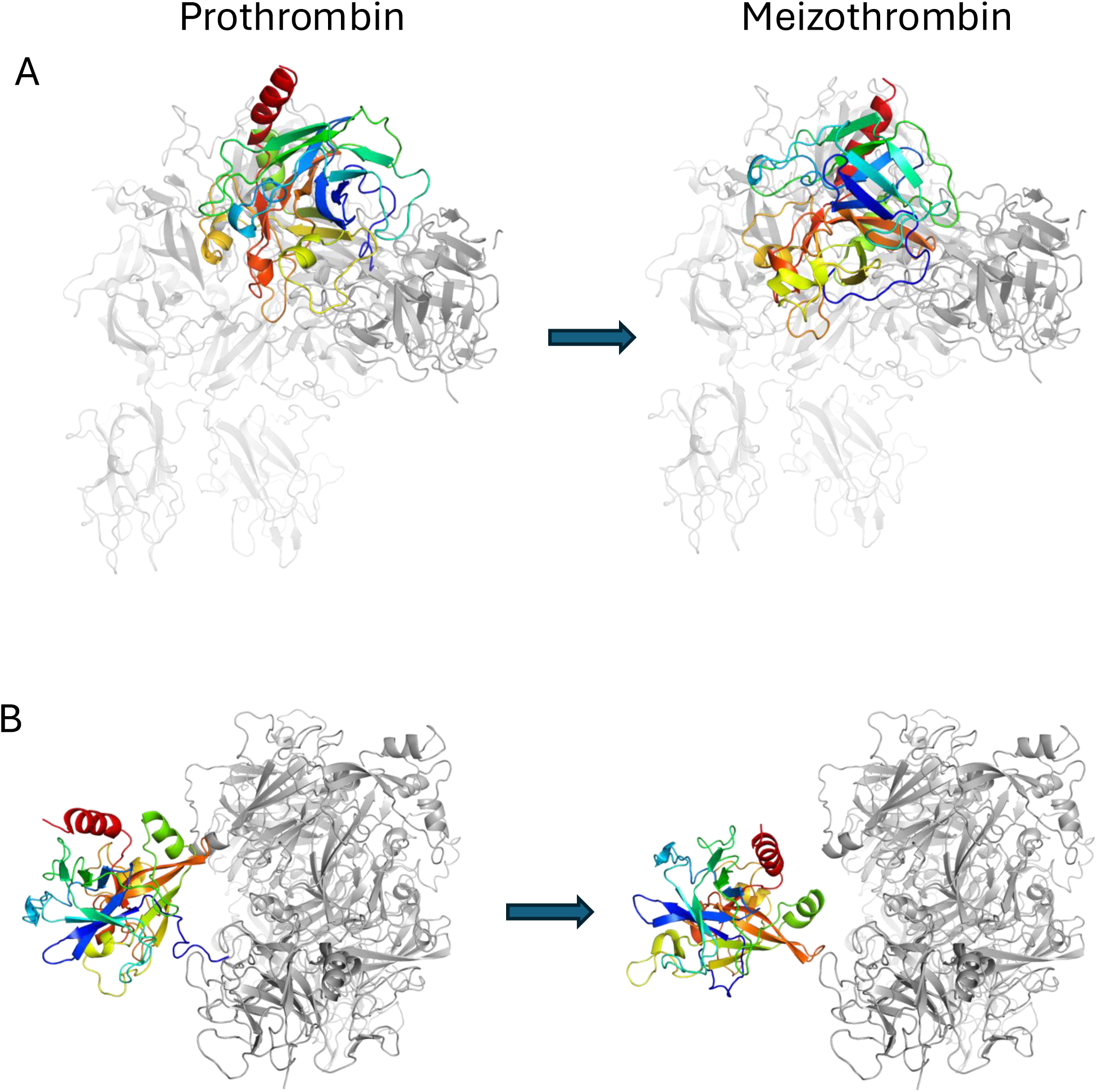
Movement and rotation of the SP domain of prothrombin upon conversion to meizothrombin. The SP domain of prothrombin (left) and meizothrombin (right) are depicted in cartoon representation, colored from N-to-C terminus (blue-to-red) to illustrate the magnitude of positional and rotational shift that occurs after cleavage of Arg320. The side view (A) and top view (B) are shown, with prothrombinase colored gray. A movie of the morph in each orientation is provided in Supplementary materials.

**Figure 6.**
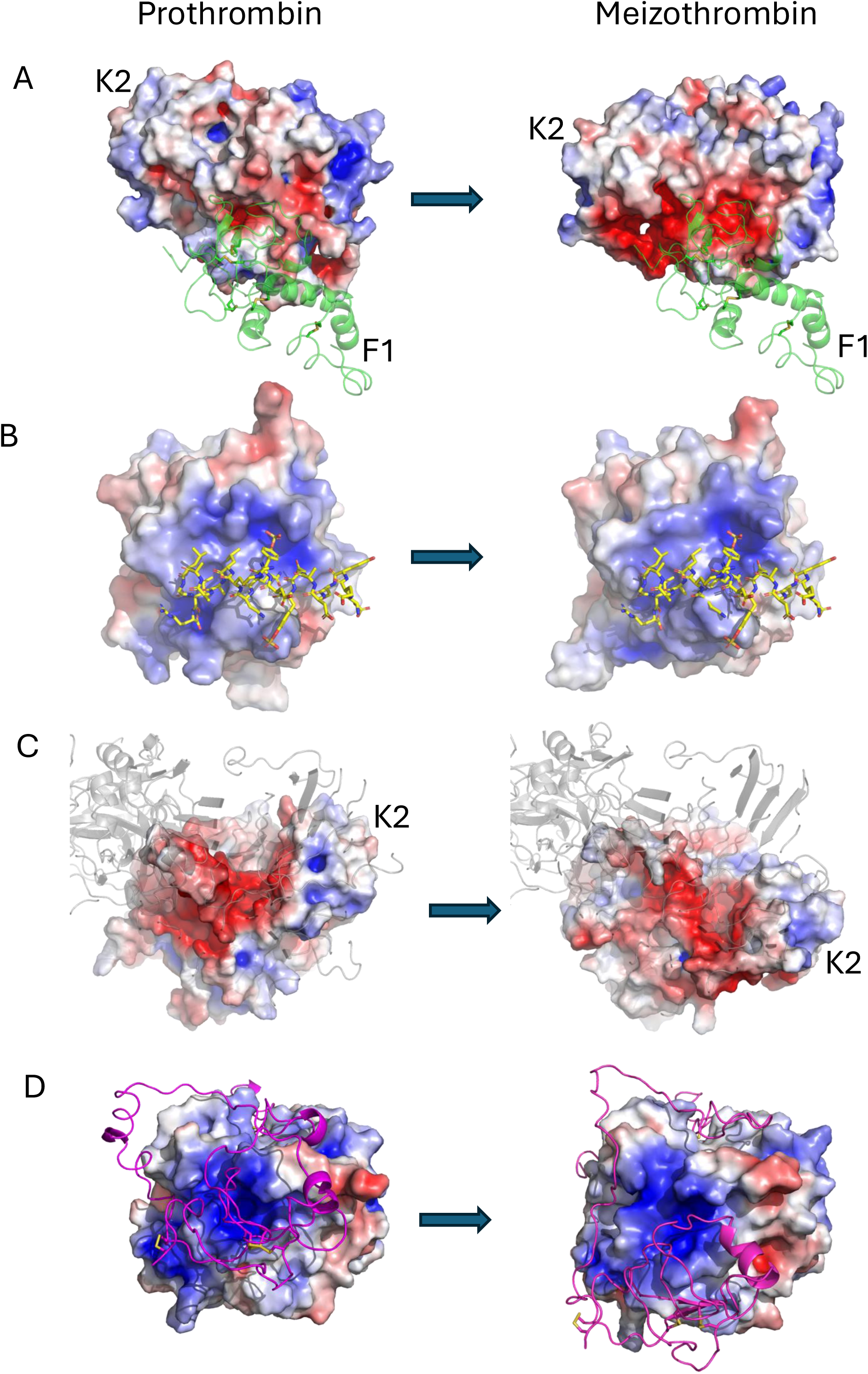
Surface elecrostatic representations of the SP and K2 domains of prothrombin (left) and meizothrombin (right) with interaction partners. (A) The F1 region of prothrombin (semitransparent green cartoon; labeled) binds in the active site of prothrombin, but is expelled upon conversion to meizothrombin (F1 is still depicted). Major changes to the shape and electrostatic properties are evident, as is movement of the K2 domain (labeled). (B) The surface of exosite I is depicted with the a2-loop in yellow sticks. Changes to the properties of exosite I are evident upon conversion to meizothrombin (the a2-loop is still depicted). (C) The surface of prothrombin and meizothrombin presented to fVa (gray cartoon), with the SP domains fixed, illustrates the dramatic shift in shape and properties upon cleavage of Arg320. The K2 domains are indicated. (D) The SP domains are oriented identically with electrostatic sufaces shown. The K2 domain and the loop connecting it to the SP domain are depicted as magenta cartoon. Exosite II is facing and the movement of K2 along the SP domain is evident.

These active site rearrangements are common to all serine proteases upon activation, so loss of the active site interaction with F1 is easily rationalized. In contrast, although there are changes evident in exosite I (Fig. 6B), it is not clear why the interaction with the a2-loop is no longer observed. However, exosite I is known to be sensitive to the zymogen-to-protease conformational change and has been described as a pro-exosite due to the inability of prothrombin to bind to exosite I ligands of thrombin^19,20^, such as thrombomodulin^21^ and fibrinogen^22^.

Of particular relevance for the repositioning of meizothrombin to present Arg271 for cleavage is the movement of the K2 domain to alter the properties of the surface facing fVa (Fig. 6C). The K2 domain shifts 12 Å relative to the SP domain (Fig. 6D and Supp. Movie S3), resulting in a change in shape and electrostatics (Fig. 6C). In prothrombin, K2 interacts primarily with the C-terminal helix of the SP domain, centered on Lys572 (Lys240 in chymotrypsin numbering) (Supp. Table S14), but shifts in meizothrombin to interact primarily with the 90-loop centered on Arg93 (409 in prothrombin numbering; Supp. Table S15). Changes in the surface properties of exosite II upon activation of prothrombin to meizothrombin are evident (Fig. 6D) and some clashes with K2 are engendered (Supp Table S16), which together appear to drive the repositioning of K2. One other conformational change of note in conversion from prothrombin to meizothrombin is the formation of a helix in the loop N-terminal to Arg320 (Supp. Fig. S10). This new helix associates with the SP domain of meizothrombin and interacts with the loop containing Arg150 of fXa, constituting an important exosite interaction (Supp. Table S10).

## DISCUSSION

Cryo-EM captures a snapshot of a biological system, with different classes of complexes representing the distribution of states at the time of vitrification. It is interesting to note, that although we observed several possible classes for the ternary prothrombin complex, they reflected only a small degree of rotational freedom pivoting on Arg320 in the active site cleft of fXa (Supp. Fig. S3); no observed particle class presented Arg271 for cleavage. This supports exclusive processing of prothrombin at Arg320 by assembled prothrombinase^23^, and suggests that evidence of initial cleavage at Arg271 in vitro is likely artifactual and due to the fraction of free fXa. The apparent rotational freedom of prothrombin bound to prothrombinase also has important functional implications. The light-touch interaction between prothrombin and prothrombinase is unlike a normal stable protein-protein complex, with prothrombin appearing to be trapped in an electrostatic cage that enforces the presentation of the 320 cleavage site. Prothrombin is free to move about in the cage so long as the orientation is roughly the same and Arg320 is engaged in the active site of fXa. Complementary electrostatics are known to accelerate substrate association^24,25^ and help explain the cofactor effect of fVa, and, importantly, the electrostatic dependance and consequent rotational freedom allows prothrombin to undergo profound conformational rearrangements after the first cleavage event to present Arg271 without dissociation. A substrate binding mode where rigid domains interact with high affinity would inhibit/slow conformational change, and would require dissociation and reassociation to present the second binding site. Such a scenario would slow turnover and risk the loss of processivity. A loosely held substrate is also consistent with the rapid product dissociation necessary to achieve high turnover.

Now that we have determined the structures of prothrombinase in all three states, apo, substrate-bound and intermediate-bound, we are able to piece together the mechanistic detail of the processive processing of prothrombin, previously described as ‘ratchetting’^26^. Under physiological conditions, the components of prothrombinase will assemble on an activated cell, such as a platelet, that expresses phosphatidylserine on its surface. The two membrane binding C domains of fVa and the Gla domain of fXa align in a linear manner. Prothrombinase is therefore able to pivot forwards or backwards to some degree, and need not bind to the membrane surface in a strictly perpendicular orientation. However, fluorescence resonance energy transfer studies produced distances consistent with an average apo prothrombinase orientation roughly perpendicular to the membrane surface^27,28^. The situation changes when prothrombin binds to prothrombinase. The closed conformation of prothrombin predominates, and we only observe the closed configuration in our cryo-EM map, with no evidence of a smaller volume corresponding to prothrombin in any of the particle subsets (i.e. if F1 were missing). Due to the compact state of prothrombin, prothrombinase must tilt forward by about 20° to engage a prothrombin molecule bound to the same membrane surface (Supp. Fig. S11). The first state of the catalytic cycle (Fig. 7) is therefore represented by the prothrombinase-prothrombin cryo-EM structure bound to a PL surface, with prothrombinase leaning forward at an angle of about 64° relative to the PL plane (Fig. 7A). Cleavage of the Arg320-Ile321 bond then initiates a series of changes that begin with the expulsion of the new N-terminus from the active site of fXa. The non-prime side (IEGR320; P4-P1; nomenclature of Schechter and Berger^29^) should remain in place since it interacts more strongly with fXa than the prime side (321IV324; P1′-P2′).

**Figure 7.**
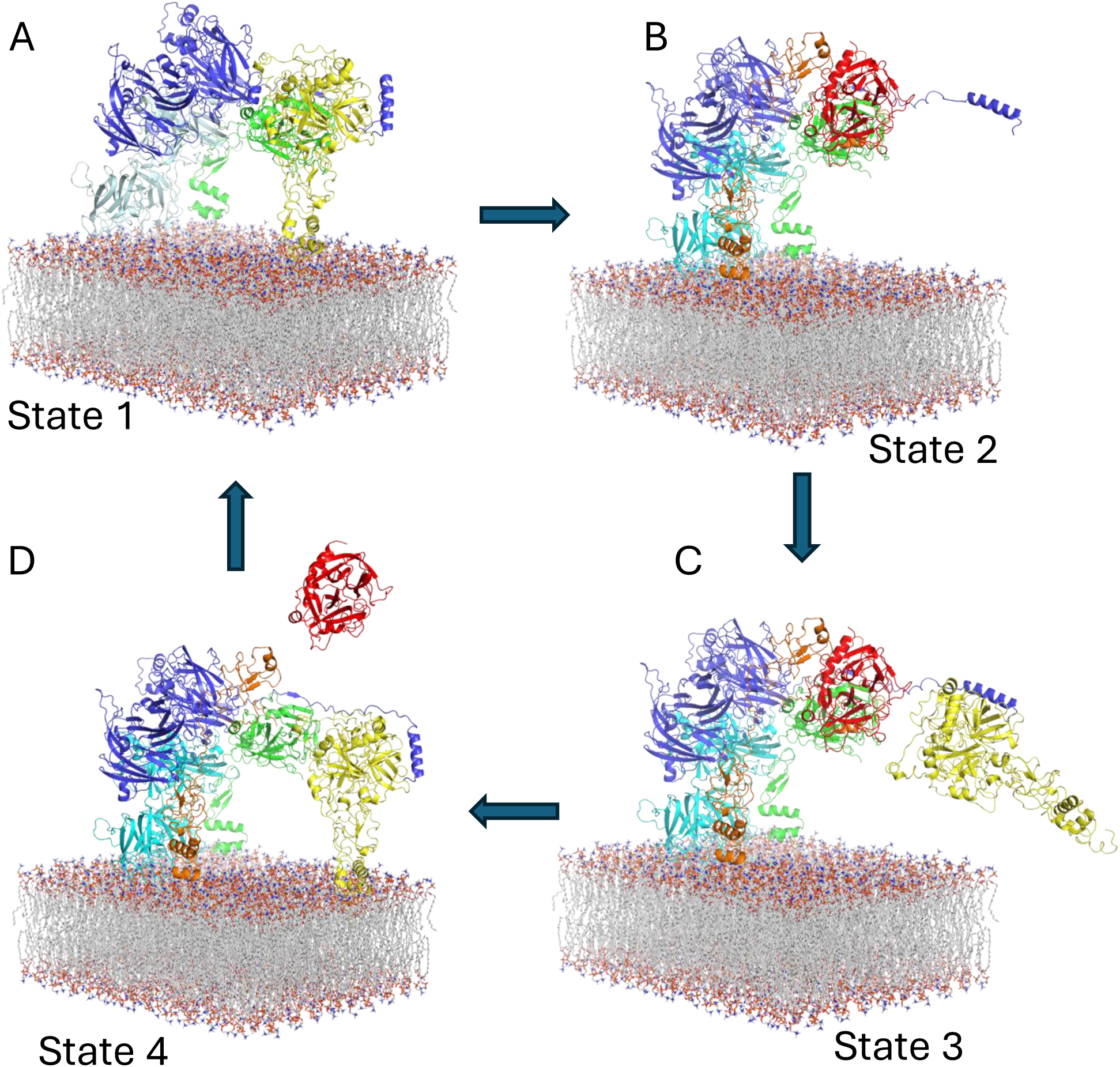
The four states of the catalytic cycle of prothrombin processing by prothrombinase. The prothrombinase complex bound to a phospholipid (PL) bilayer is depicted, as in Fig. 1, with the Gla and EGF1 domains of fXa from the apo structure added. (A) Prothrombin (yellow) bound to prothrombinase in a productive complex, with all three proteins binding the PL surface (State 1). Prothrombinase is oriented at an angle of about 64° relative to the plane of the PL membrane. (B) State 2 following cleavage at Arg320 to form the meizothrombin intermediate (orange for F1.2 and red for active SP domain). F1.2 has moved to accommodate the rotation of the SP domain and the C-terminal portion of the a2-loop has disengaged. Prothrombinase relaxes back to a near perpendicular orientation with respect to the PL surface. (C) We hypothesize that before or during cleavage of Arg271 that the a2-loop will engage another prothrombin molecule from either the PL-bound or free (depicted) pool (State 3). The Kd of prothrombin for an activated PL surface is similar to its concentration in blood, so both pools are likely to be equally populated^52^. (D) Cleavage of Arg271 expells the product thrombin (red; State 4) and prothrombin can now move into the embrace of prothrombinase, attracted by long-range electrostatics, and the remnant F1.2 is free to diffuse away, returning to State 1 and completing the cycle.

Once the new N-terminus is free, it becomes a tethered ligand and rapidly finds its way to the center of the SP domain of prothrombin. The zymogen-to-protease conformational change then occurs to the SP domain^17^, which, due to the allosteric linkage between the active site, exosite I and exosite II of prothrombin/thrombin^19–21,30–32^, all contacts involving the SP domain are dramatically altered. The following rearrangements are likely to be simultaneous: dissociation of F1 from the SP domain; dissociation of the a2-loop of fVa from exosite I; and, shifting of the K2 domain on exosite II of the SP domain. We propose that the K2 domain shift triggers the 50° rotation of the SP domain relative to prothrombinase, and that this rotation extracts the C-terminus of the light chain of meizothrombin (317IEGR320) from the active site of fXa. The extracted C-terminus then forms a helix to mediate a new exosite contact with the 150-loop of fXa. The rotation of the K2/SP unit also, importantly, places Arg271 into the active site of fXa. The release of the F1 region from the SP domain of meizothrombin appears to be necessary to allow the rotation of the SP domain whist maintaining Gla domain contact with the PL (Supp. Fig. S12). Once dissociated from the SP domain, F1 will also need to shift towards the C2 domain of fVa in order to remain bound to the PL surface, due to the limited length of the linker between K1 and K2. Elongation of the linker and movement of F1 towards fVa will allow prothrombinase to tilt backwards to obtain a more perpendicular orientation with respect to the PL surface. This PL-bound form of the meizothrombin-prothrombinase complex represents State 2 in the catalytic cycle (Fig. 7B).

In contrast, dissociation of the a2-loop from meizothrombin is not required to allow the rotation of the SP domain, with the linker between the N- and C-terminal acidic regions being of sufficient length to accommodate the shift in exosite I position (Supp. Fig. S13). We propose that its dissociation instead plays a role in potentiating high turnover. The first obvious effect of losing the a2-loop contact is that meizothrombin is held less firmly in place and will dissociate faster after cleavage of Arg271. The second and less obvious effect is that release of the C-terminus of the a2-loop allows it to bind to a prothrombin molecule before dissociation of fully processed thrombin. This pre-loading of prothrombin from either the PL-bound or free pools would increase the rate of turnover and constitutes State 3 in the catalytic cycle (Fig. 7C).

Meizothrombin is thus poised to dissociate as thrombin once cleavage at Arg271 occurs. As for the first cleavage event, the principal contacts in the active site of fXa are on the non-primed side. The prime side, now constituting the light chain of thrombin, will be expelled once cleavage has occurred. F1.2, comprised of F1 and the K2 domain, is now only attached to thrombin by non-covalent association with exosite II, and in the absence of tethering, this interaction is very weak with a dissociation constant of 3.7 μM^33^. Thrombin is therefore free to dissociate leaving the C-terminus of F1.2 still in the active site of fXa, constituting State 4 of the catalytic cycle (Fig. 7D). F1.2 will then diffuse away as the pre-captured prothrombin docks to begin another round of the catalytic cycle.

Another implication of this work relates to the subject of thrombin allostery. Thrombin has long been considered an allosteric enzyme, with cross talk between the active site, the Na^+^ binding site, exosite I and exosite II^34^. However, there is no evidence that allostery is actually important for any of the multiple activities of thrombin, including the most radical switch in activity from pro- to anti-coagulant upon thrombomodulin binding to exosite I. This and other activity changes in thrombin rely on exosite competition and not on allosteric switching^35^. What is abundantly clear from the structures described in this manuscript is that allostery plays an essential role in prothrombin processing. The apparent allosteric connections between the active site and exosites on thrombin may simply be the remnants of the allostery required for sequential and rapid processing of prothrombin, and be of no further functional relevance.

Finally, it should be noted that two cryo-EM structures purporting to be of the prothrombinase-prothrombin complex have been deposited^36,37^. We have previously detailed the shortcomings of 7TPP^38^, which is a low-resolution structure (likely 5.5 Å [<u>PubPeer</u>]) from grids made in the absence of PL. The deposition 7TPQ, originally of apo prothrombinase formed on nanodiscs, has been replaced with 9CTH which claims to be of the prothrombin-prothrombinase complex on nanodiscs at 6.5Å resolution^37^. However, careful analysis has demonstrated that the prothrombin component is entirely missing from the map, and likely represents a case of Einstein-from-noise^39^ (<u>PubPeer</u>). We conclude that the structures presented in this manuscript are the first of sufficient quality and resolution to decipher the complex and surprising mechanism by which prothrombin actively participates in its own processing, a mechanism we term ‘substrate allostery’.

## METHODS

### Recombinant protein expression and purification

B-domainless fV was expressed, purified and activated as previously^40^. The minimal M17 fXa construct containing only the EGF2 and SP domains was used in place of full-length protein because the Gla and EGF1 domains contribute little to fVa binding^12^ and were not well defined in the map of our previous cryo-EM structure^13^. The S195A variant of M17 fX was produced in E. coli as inclusion bodies, refolded, purified and activated as previously described^23^. Prothrombin (S195A) was prepared as described previously^41^. Meizothrombin was generated from S195A prothrombin as previously^42^. The purity and integrity of meizothrombin was confirmed by SDS-PAGE.

### Cryo-EM grid preparation, data collection, processing and refinement

Grid preparation and data collection were conducted essentially as previously^13^. Briefly, fVa was mixed with truncated S195A fXa (M17 variant) and S195A prothrombin or meizothrombin at a 1:6:2 molar ratio, with final concentrations of 650 nM, 3.9 μM and 1.3 μM, respectively. All proteins were in 20 mM HEPES pH 7.5, 150 mM NaCl and 5 mM CaCl2. Gold grids (Quantifoil (AU) R1/1 on Au 300 mesh) were glow discharged in residual air at 25A for 60 s using a Pelco EasiGlow, and 3 µl of protein complex solution was applied. Following a 10 s wait time, grids were blotted for 1 s using -7 (prothrombin) or 0 (meizothrombin) blot force in a 100% humidity chamber at 4°C, and then flash plunged into liquid ethane (-180°C) using a Vitrobot Mark IV. The prothrombin complex was imaged using a Titan Krios electron microscope with a K3 detector (Gatan) at the Cambridge Nanoscience Centre. Data were collected at 130,000 magnification (pixel size 0.829 Å) at a dose of 50 e−/Å^2^ and defocus values between -1.8 and -0.6 μm. Grids of the meizothrombin complex were imaged using a Titan Krios with a Falcon4i counting camera at the University of Cambridge, Department of Biochemistry Cryo-EM facility. Data were collected at 165,000 magnification (pixel size 0.729 Å) at a dose of 50.3 e− /Å^2^ and defocus values between -1.8 and 0.8 μm. All data were collected on a tilted stage (-25°) to improve the particle orientation distribution.

All data were processed in cryoSPARC^43^. Patch motion correction and patch CTF were first run using default parameters; exposures with an estimated CTF resolution fit higher than 4.6 Å or a defocus tilt angle larger than 30 deg were discarded. For the prothrombin complex, blob picker was used to pick and extract 7,138,515 particles that were extracted in 320x320 pixel boxes. Several rounds of ab initio reconstructions, heterogeneous refinements, and 3D classifications were used to filter out bad particles and generate a ‘consensus’ reconstruction by homogeneous refinement based on a subset of 440,004 particles representing an intact fVa-fXa-prothrombin complex. These particles were re-extracted in 384x384 pixel boxes. A single round of masked 3D classification was run to sort particles depending on the orientation of prothrombin relative to the prothrombinase complex. The mask was generated from the ‘consensus’ reconstruction and encompassed prothrombin as well as small parts of fXa and fVa using the following custom parameters: lowpass filter of 12 Å, threshold of 0.02, and soft padding width of 18 pixels. Five distinct classes were obtained (Supp. Fig. S2). The best defined map was further refined using non-uniform refinement to a final resolution of 3.1 Å using 76,039 particles (Supp. Fig S3A). The ‘consensus’ subset was also used to obtain a map at higher resolution for prothrombin by masked local refinement. The mask described above was used again, along with a custom parameter of rotation search extent of 20 degrees, to generate a 3.4 Å map. A composite map was constructed using the original map focused on prothrombinase and the one focused on prothrombin. Briefly, most of the map corresponding to prothrombin was removed from the best overall map using ChimeraX^44^ and sharpened in Phenix^45^. Similarly, map corresponding to prothrombinase was removed from the prothrombin-focused map and then sharpened. A composite map was then made in Phenix. This the resulting composite map preserved the best features of each and was used for model building. For the meizothrombin complex, a final set of 25,376 particles was similarly re-extracted to generate a final map of 3.1 Å resolution (Supp. Fig. S2B). A similar issue of rotational freedom of meizothrombin was observed in the map (Supp. Fig. S4), and a composite map was made for model building, as before.

The M17-prothrombinase structure (9I2H) was fit into the map of the prothrombin complex in ChimeraX^44^. The AlphaFold^16^ model of prothrombin (AF-P00734-F1-v6) was chosen as a starting model for prothrombin, due to the lack of any high-quality crystal structure and the fact that it fit well into the cryo-EM map. The first 43 residues were removed and the mature protein was renumbered. Gamma carboxylation was not performed to the coordinates because the map was of insufficient quality in that region to place side chains. The model of prothrombin was fit into the map using ChimeraX. The two C domains were subsequently subjected to rigid body refinement in Coot^46^. Residues corresponding to F1 of prothrombin were also subjected to rigid body refinement in Coot, and underwent a small shift relative to starting model. Similarly, 9I2H was placed into the map of the meizothrombin complex in ChimeraX. It was immediately clear that the volume corresponding to meizothrombin did not contain the F1 region, and fit well to the crystal structure of desF1 meizothrombin 3E6P^47^ which was placed in the map using ChimeraX. Rigid body refinement was conducted in Phenix^45^ for each chain, and for the two C domains of fVa. Model building was conducted using Coot and refinement with Phenix. Data collection, processing and refinement statistics are provided in Supp. Tables S1 and S2.

### The Pre-2-a2-peptide crystal structure

An N-terminally acetylated peptide corresponding to the C-terminal region of the a2-loop (EPEDEESDADY*DY*QNRLAAALGIR; * indicates phosphorylation; Biomatik) was dissolved in 10 mM Tris pH 8.0 to a final concentration of 10 mM. A 5-fold molar excess of peptide was added to a solution containing 10 mg/ml Pre-2 to form complex prior to crystallization. Crystallization was performed by sitting drop vapor diffusion in 96-well MRC crystallization plates (SWISSCI) using the Mosquito LCP (TTP Labtech) to dispense 100 nl of protein and 100 nl of reservoir solution. Crystallization was monitored using a Rock Imager 182 (Formulatrix). Crystals of various morphologies were obtained in several conditions and were cryoprotected with 20% glycerol, flash-cooled in liquid nitrogen and sent to Diamond Light Source (Oxford, UK) beamline I03 for data collection. The best dataset was obtained from a crystal grown in Index screen (Hampton Research; 0.1 M Tris pH 8.5, 0.2 M NaCl, 25% PEG3350). Data processing was conducted using AIMLESS^48^ and molecular replacement was conducted in Phaser^49^ with an AlphaFold^16^ model of the Pre-2-peptide complex. Model building and refinement were conducted in Coot^46^ and Refmac^50^, and structure validation was conducted with MolProbity^51^. Processing and refinement statistics are given in Supp. Table S3.

## Acknowledgements

This work was part funded by British Heart Foundation Programme (RG/16/9/32391) and Project (PG/24/11721) grants to JAH, and Cancer Research UK Discovery Award (DRCNPG-Jun24/100002) and UK Medical Research Council (UKRI1443) Programme Funding to AJW. We thank Dima Chirgadze, Giulia Paris and Lee Cooper at the Department of Biochemistry Cryo-EM Facility, University of Cambridge and Sigurdur Thorkelsson, Pablo Castro Hartmann and Scott Gardner at the Cambridge Nanoscience Cryo-EM facility for their expert assistance.

## Authorship Contributions

JAH and FIU conceived of and designed the study. FIU and AF conducted the experiments. JAH, FIU, AF and AJW interpreted the data, and JAH wrote the manuscript.

