## Supplemental Information for "Prothrombinase processivity is conferred by substrate allostery"

### Supplemental Figures

**Fig. S1** (A) Schematics of prothrombin domain organization in open and closed states. The N-terminal gamma carboxyglutamic acid (Gla) domain is associated to the first kringle domain (K1), followed by a 26-residue linker, and the K2 and serine protease (SP) domains that are tightly associated. The closed form, where K1 interacts with the active site of the SP domain predominates in solution (B) Prothrombin processing pathways are shown with domains labeled as in (A). fXa on its own will cleave first at Arg271 (left) to form prothrombin-2 (Pre-2) and F1.2, and prothrombinase cleaves initially at Arg320 to activate the SP domain (red) and form meizothrombin. (C) The sequence of the a2-loop of factor fVa is given, with the two acidic regions containing sulfated tyrosines underlined. * denotes sulfation.

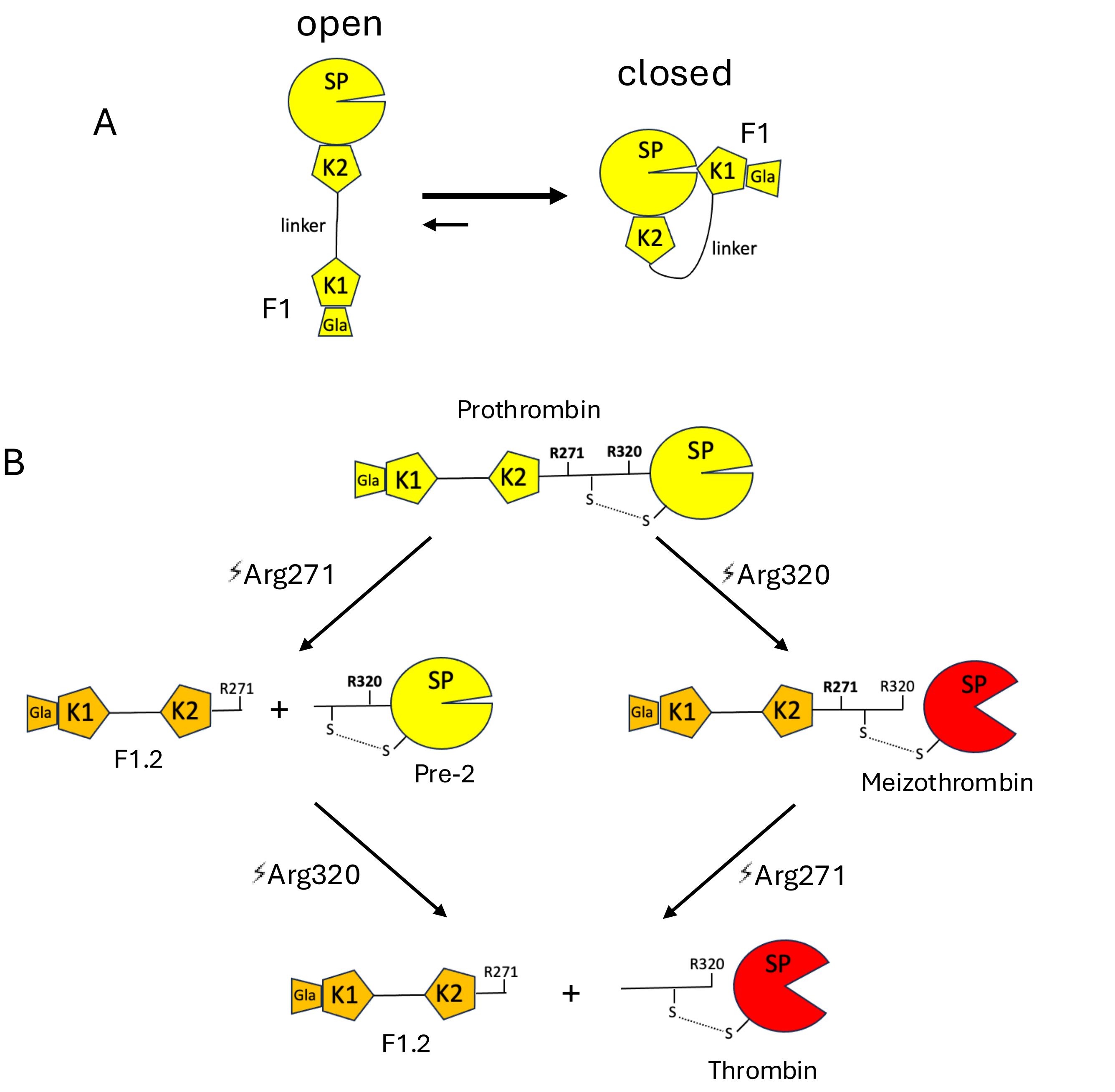

C

$_{657}$IPDDDEDSY*EIFEPPESTVMATRKMHDRLEPEDEESDADY*DY*QNRLAAALGIR$_{709}$

**Fig. S2** Fourier Shell Correlation (FSC) plots for the maps of the prothrombinase-prothrombin (A) and prothrombinase-meizothrombin (B) complexes.

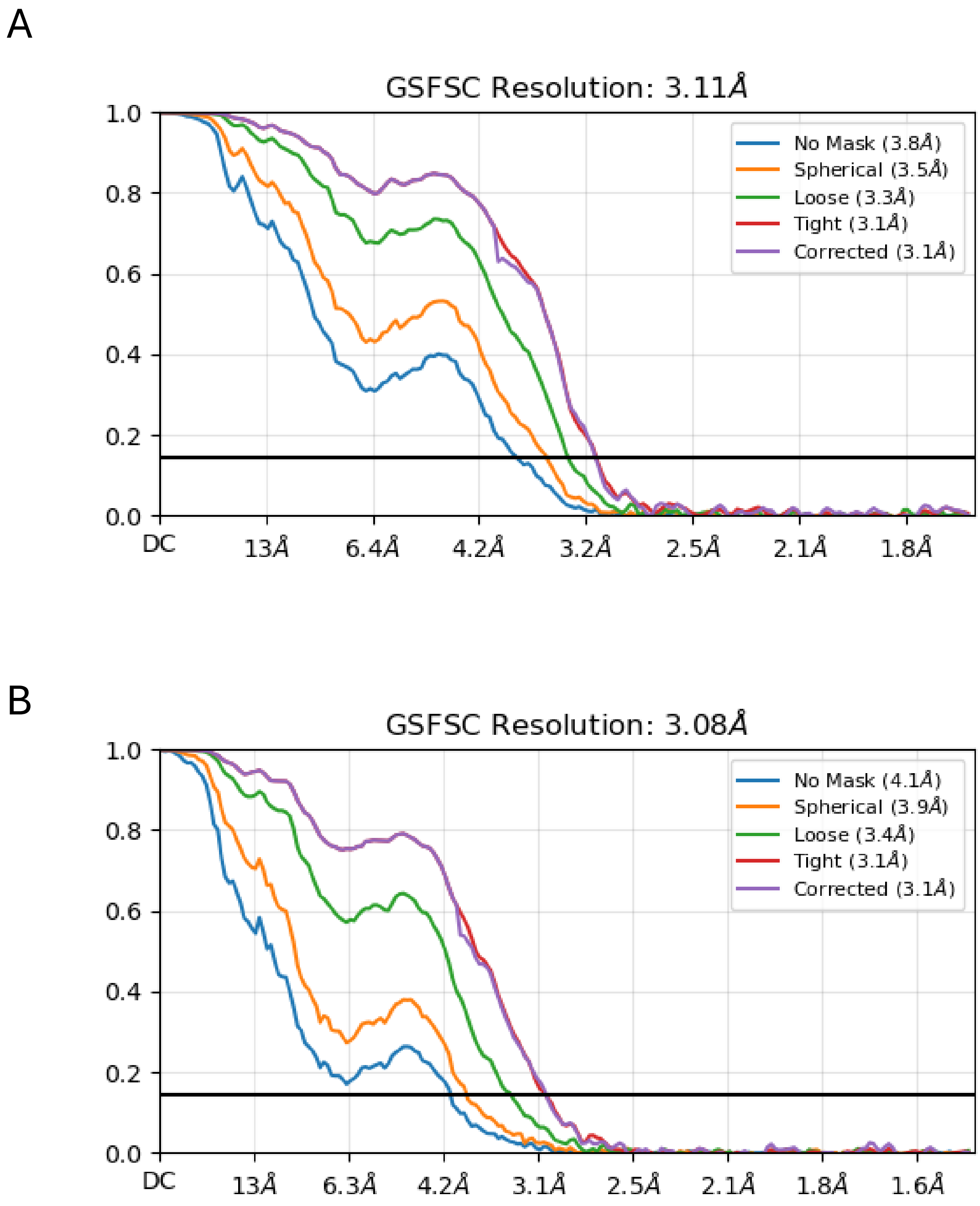

**Fig. S3** Five particle subset classes of the prothrombin component of the prothrombinase-prothrombin complex is illustrated by the fit of prothrombin into the resulting maps. Top and side views are shown. Gray is prothrombinase and the colored cartoons are prothrombin.

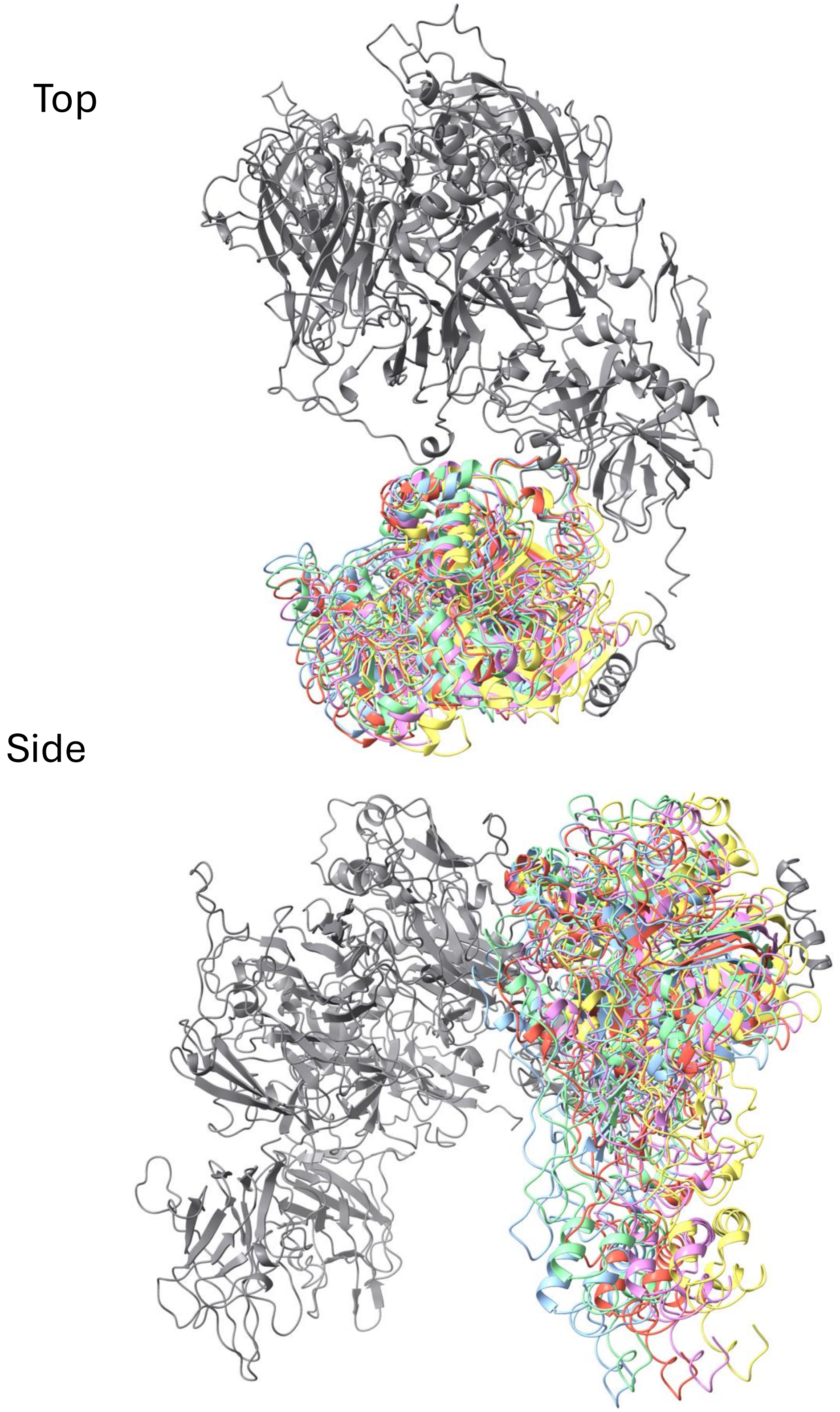

**Fig. S4** Five particle subset classes of the meizothrombin component of the prothrombinase-meizothrombin complex is illustrated by the fit of meizothrombin into the resulting maps. Top and side views are shown. Gray is prothrombinase and the colored cartoons are prothrombin.

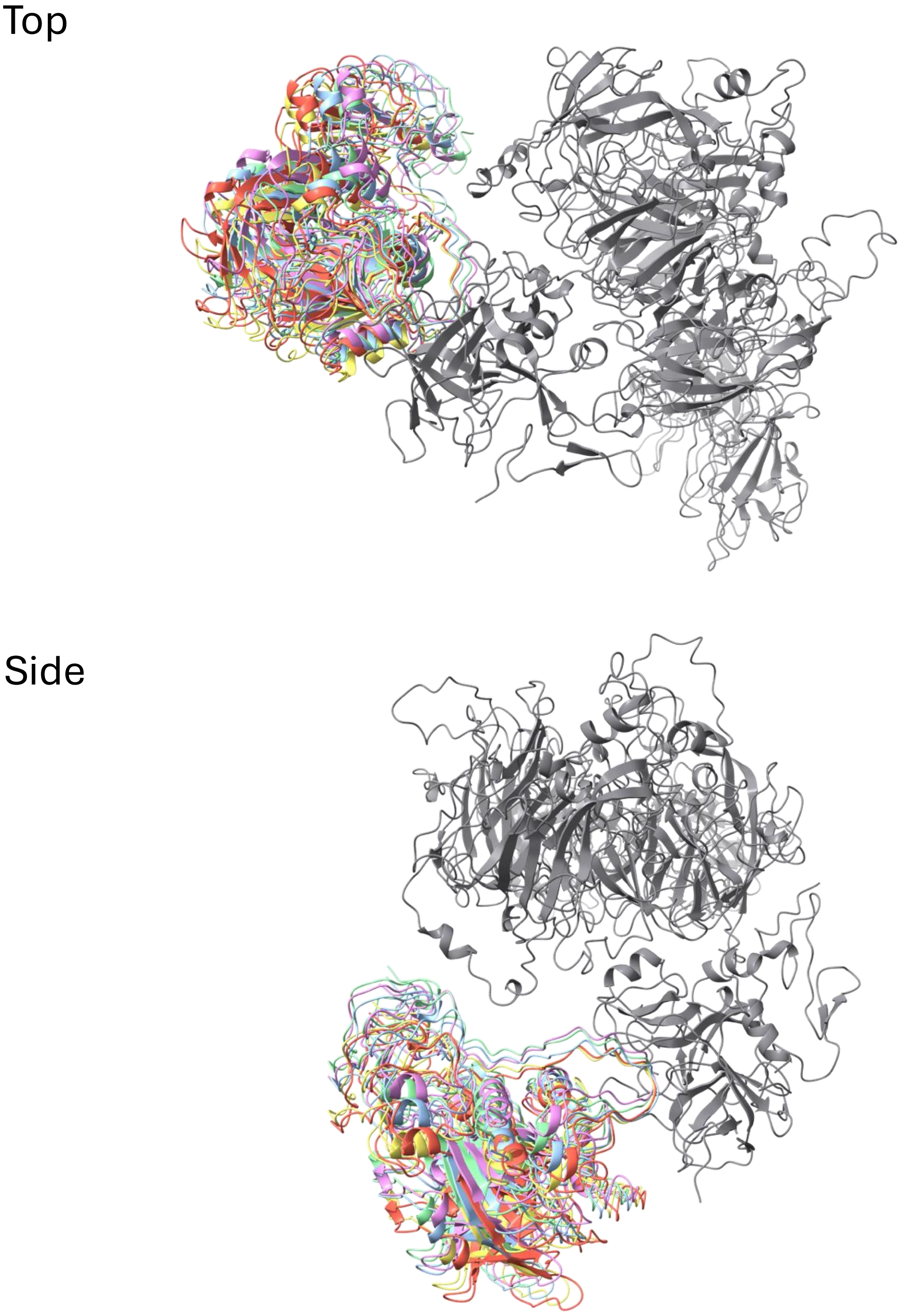

**Fig. S5** Stereo view of the active site of fXa (green) from the prothrombinase-prothrombin complex, with residues 317-322 of prothrombin (yellow sticks) surrounded by map (semitransparent gray). The figure is in the standard orientation, so that N-terminal portion of the substrate loop is on the left and the C-terminus on the right.

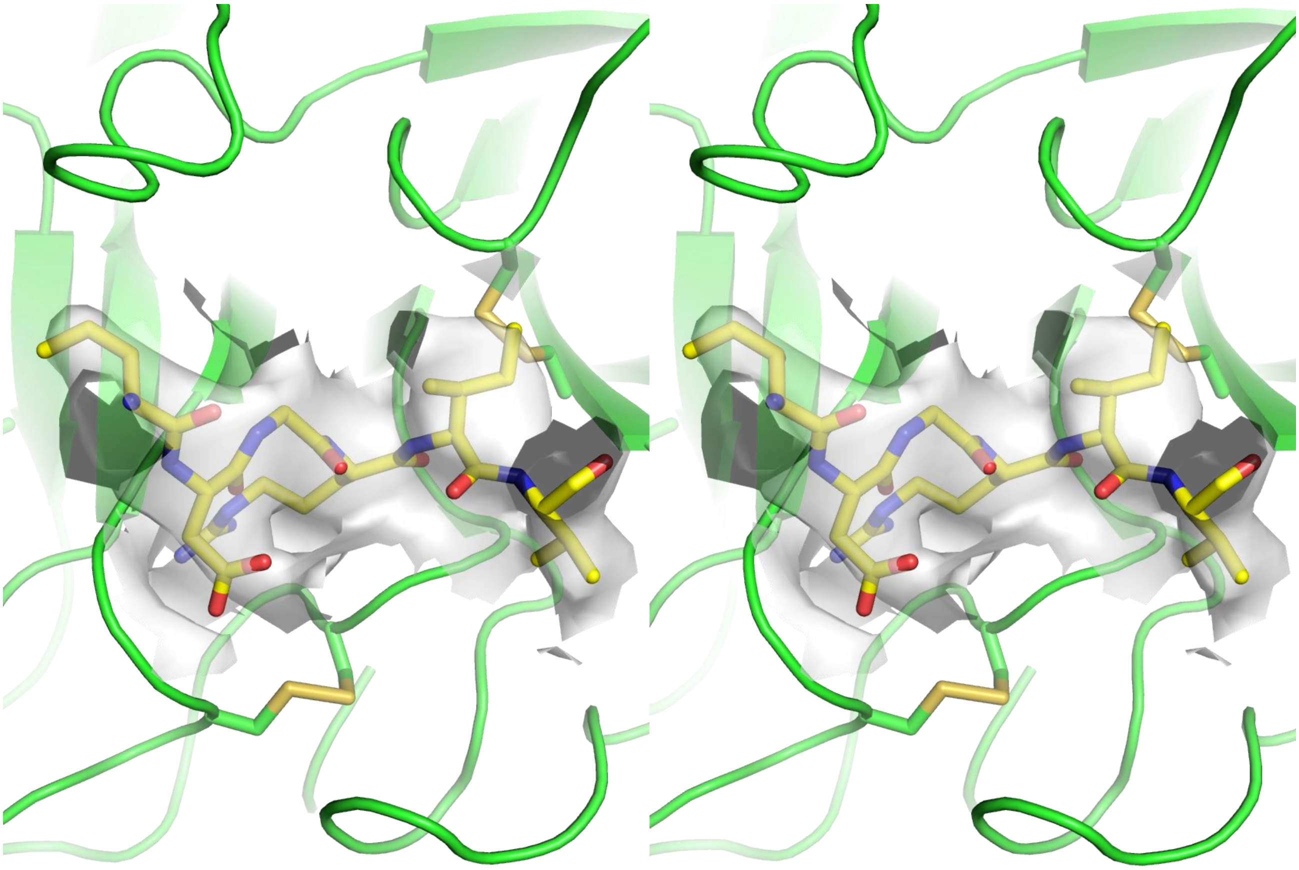

**Fig. S6** Screenshot of the AlphaFold results using the sequences of prothrombin-2 and a peptide corresponding to the C-terminal acidic region of the a2-loop. The result is colored according to the confidence score (key above), and the 2D relative positional confidence plot on the right.

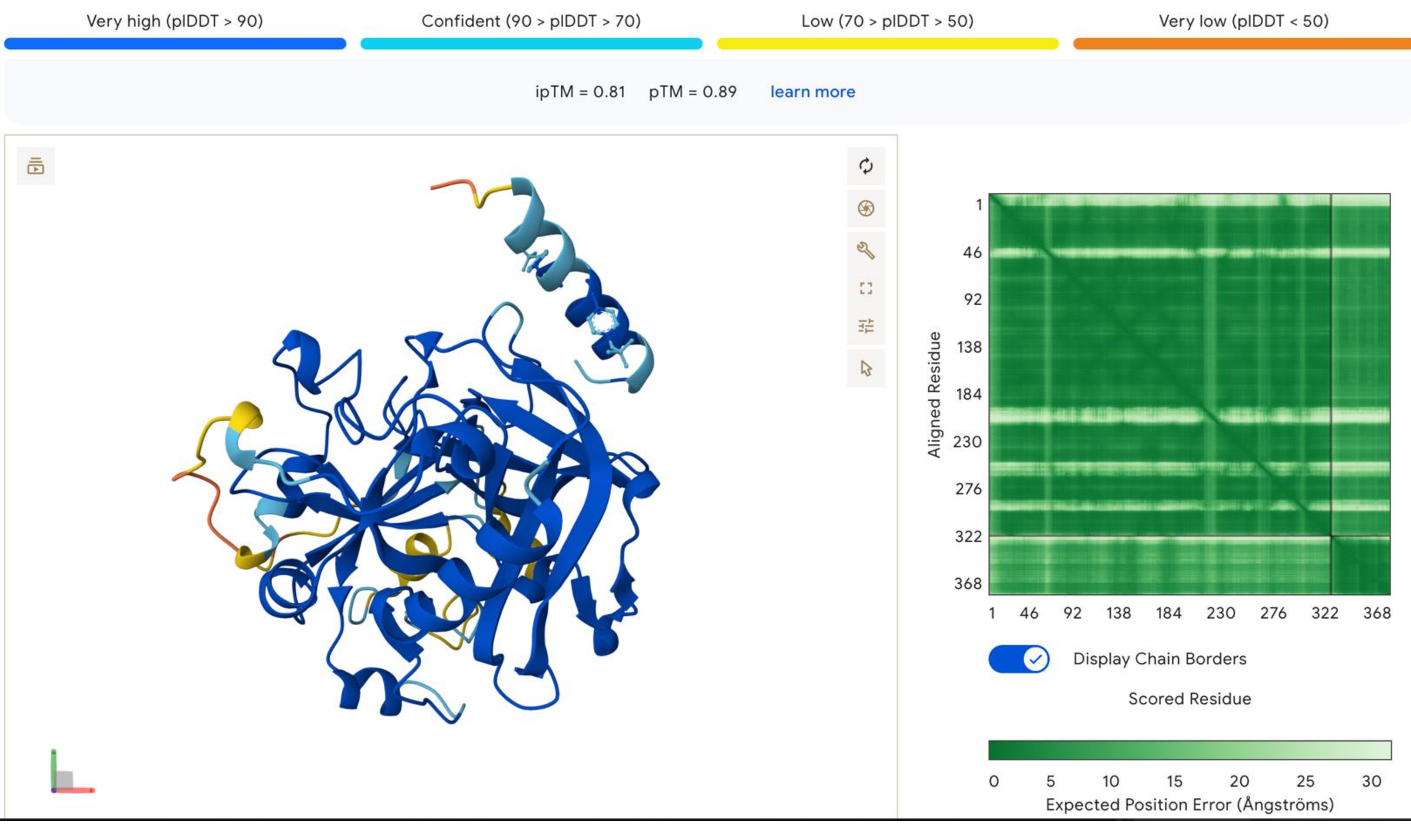

**Fig. S7.** Stereo views of the predicted and experimentally-derived interactions between prothrombin and the C-terminal portion of the a2-loop. (A) The AlphaFold models of the Pre-2 complex with an a2-loop peptide. Pre-2 is colored from N-to-C terminus (blue-to-red) and the five a2-loop predictions are in magenta, with the C-terminal Arg709 and the two phosphorylated Tyr residues depicted as sticks. (B) Electron density (blue mesh) of the crystal structure of Pre-2 (green) around the a2-loop peptide (yellow sticks). (C) The AlphaFold solution (magenta), the crystal structure (yellow), and the cryo-EM structure of the prothrombinase-prothrombin complex (blue) of the a2-loop regions are superimposed to demonstrate the overall conservation of binding interface.

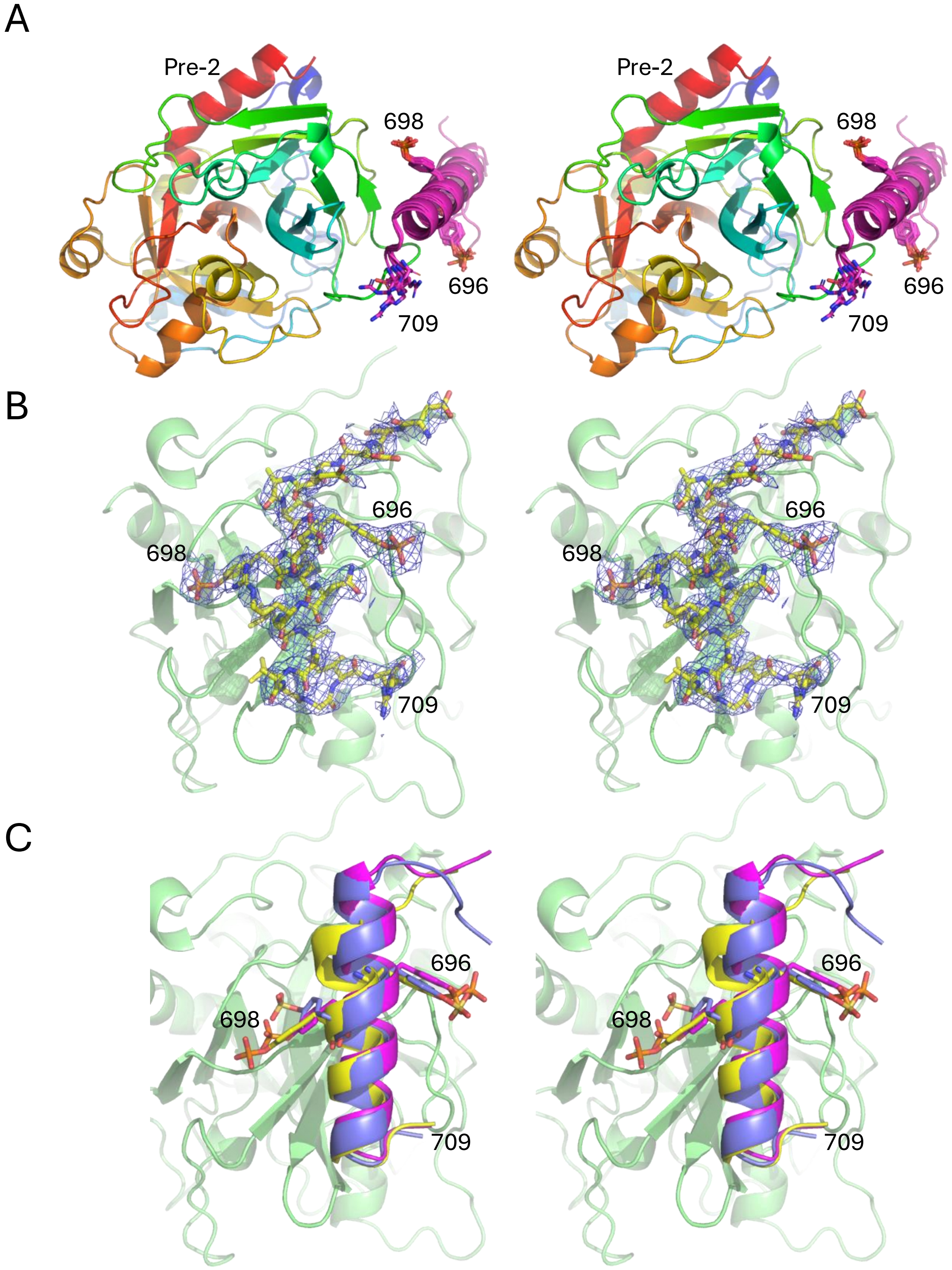

**Fig. S8** (A) Stereo view of the active site of fXa (cyan) from the prothrombinase-meizothrombin complex, with residues 268-273 of meizothrombin (orange sticks) surrounded by map (semitransparent gray). The figure is in the standard orientation, so that N-terminal portion of the substrate loop is on the left and the C-terminus on the right. (B) Superposition of the prothrombin and meizothrombin interactions in the active site of fXa, colored as in Fig. S5 and (A).

A

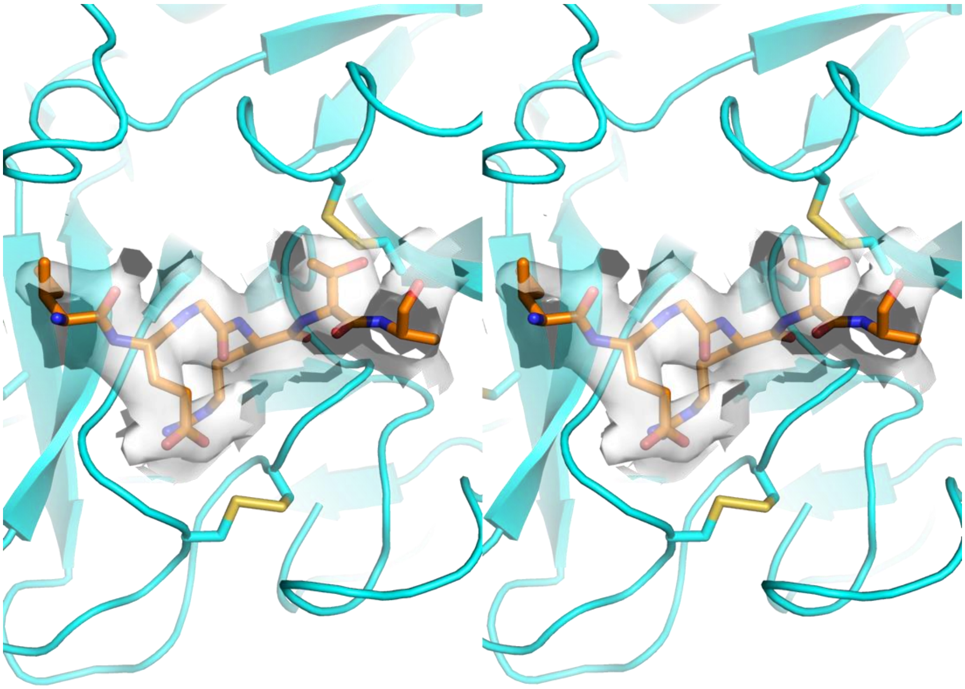

B

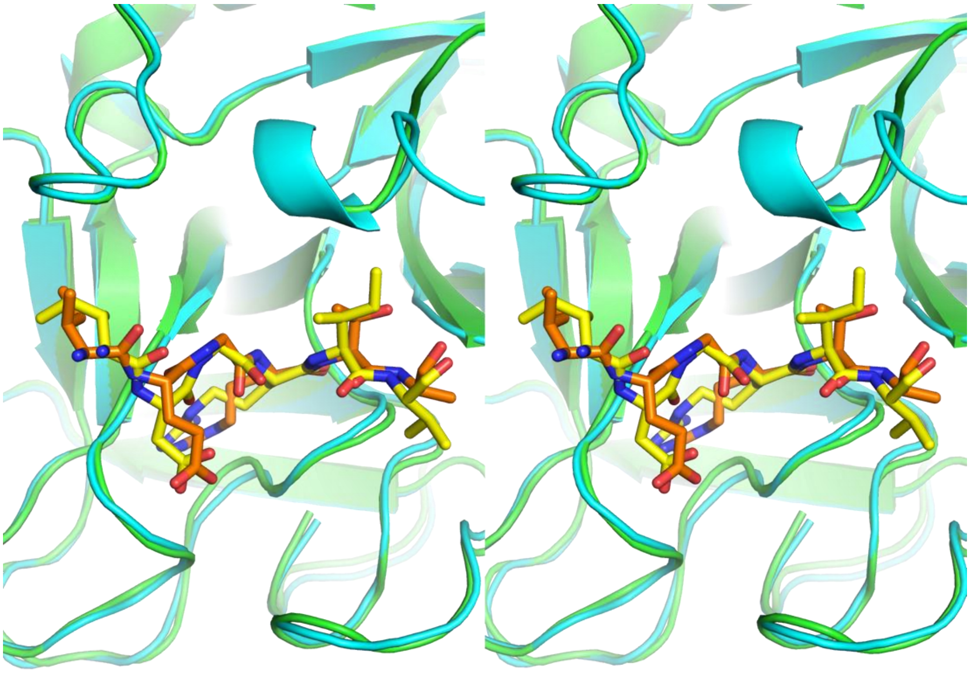

**Fig. S9** Stereo view of the superposition of the prothrombinase component of the apo (9I2H; gray with Gla-EGF1 removed), prothrombin (9TLE; yellow) and meizothrombin (9TLG; orange) structure.

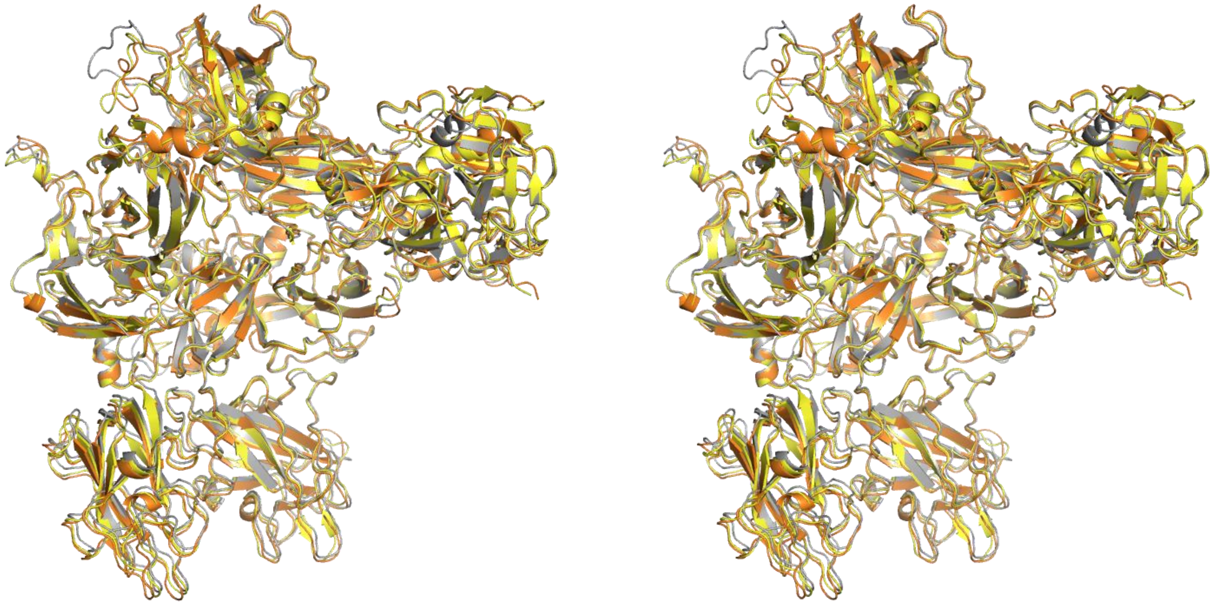

**Fig. S10** Transition of the 320-loop of prothrombin into a helix after cleavage, with only the K2-SP domains shown as cartoon, colored from N-to-C terminus (blue-to-red). The new 320-helix is indicated.

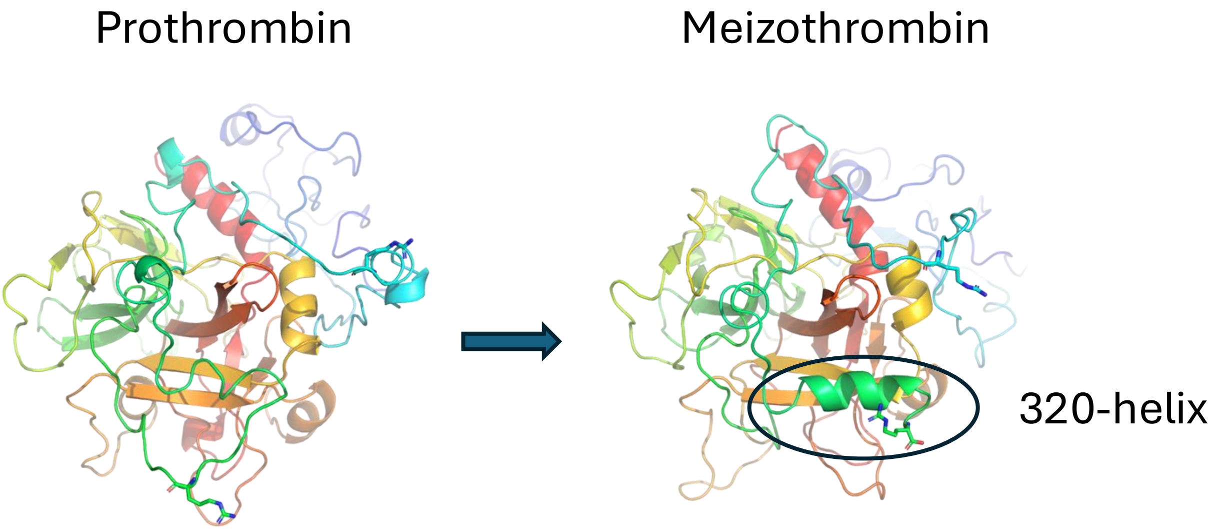

**Fig. S11** A side view of the full-length prothrombinase structure (Gla and EGF1 as in 9I2H) and a model of a PL bilayer illustrates the potential range of movement. The near perpendicular orientation of apo is shown on the left and the tilting required to accommodate PL-bound prothrombin (holo) is on the right. Roughly 20 degrees forward tilt is required.

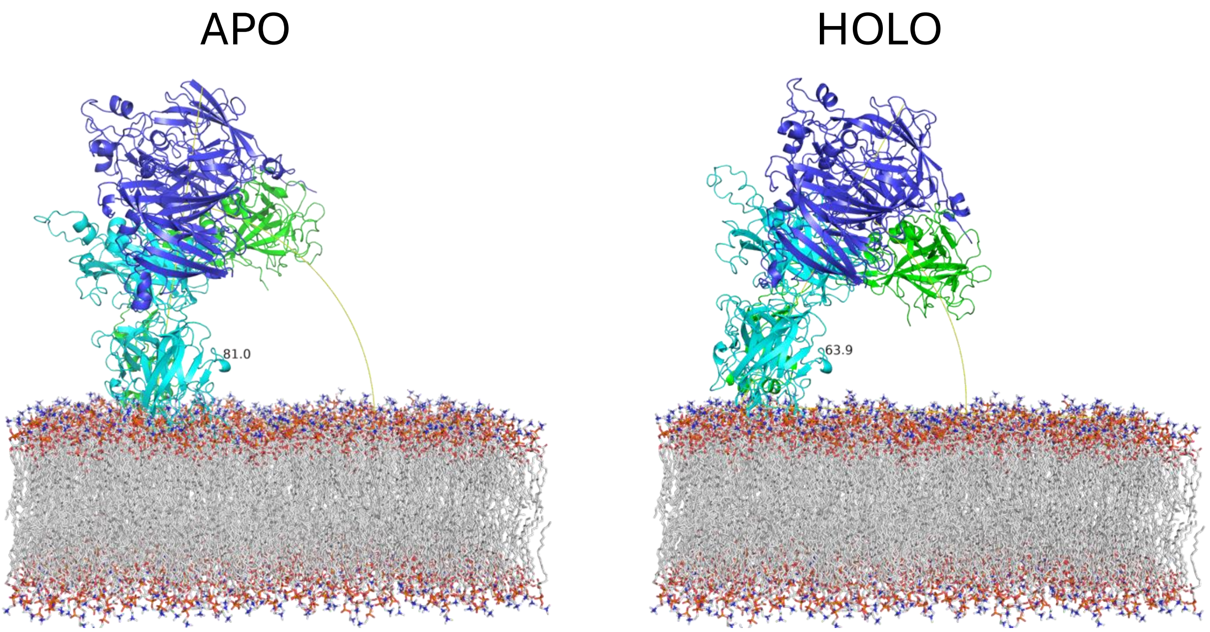

**Fig. S12** The prothrombin-to-meizothrombin transition modelled on a PL bilayer surface illustrates the movement of the F1 region (Gla and K1) if dissociation from the SP domain did not occur. Prothrombin is colored as in other figures (fVa heavy chain in blue and light chain in cyan; fXa in green; prothrombin in yellow and meizothrombin without change in F1-SP interaction in orange.

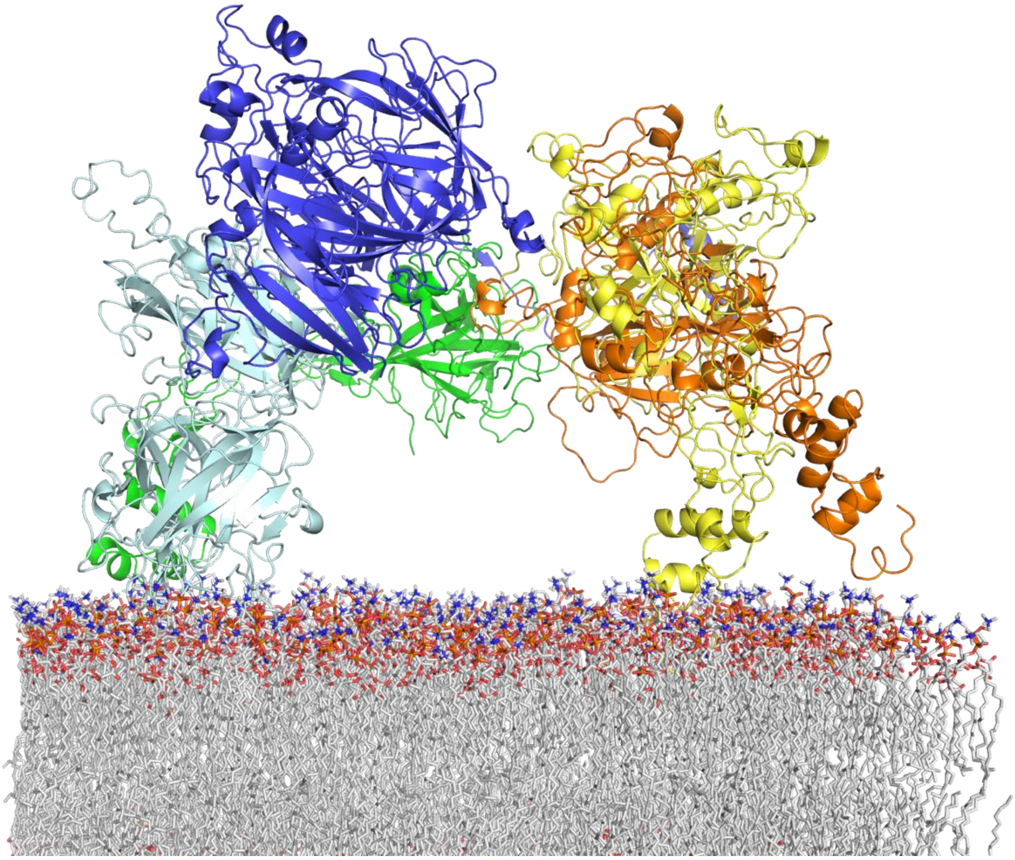

**Fig. S13** Top view of prothrombinase with meizothrombin (colored as in Fig. S12) with the C-terminal portion of the a2-loop modelled on exosite I as found in prothrombin demonstrates that the length of the linker between the two acidic portions of the a2-loop is of sufficient length to accommodate the change in SP domain orientation.

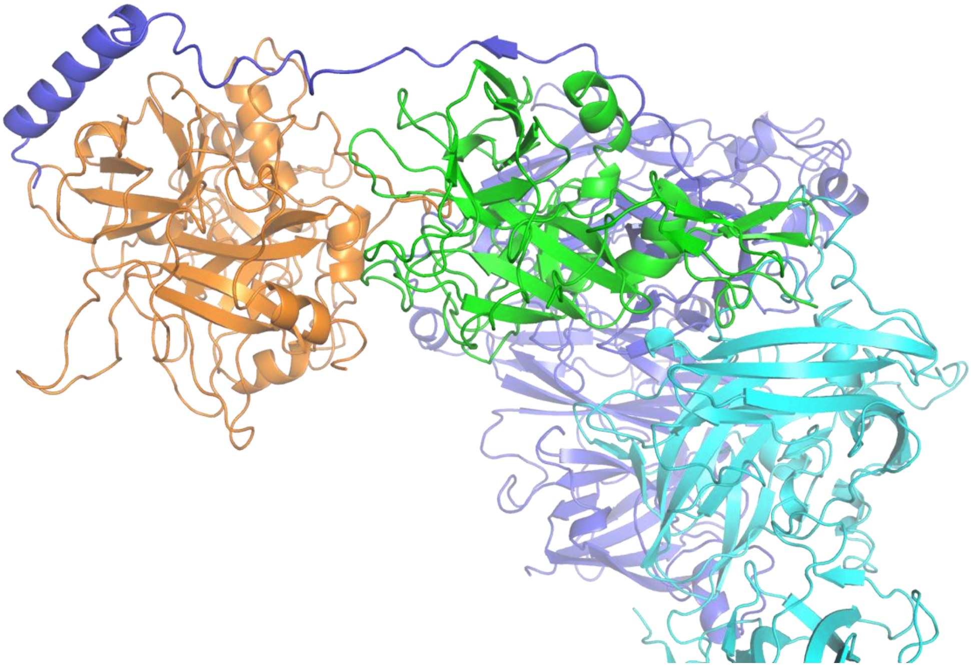

**Supplemental Table S1.** Cryo-EM data collection, refinement and validation statistics for the prothrombinase-prothrombin complex (EMD- 56052; PDBID 9TLE).

| | Prothrombinase-Prothrombin Complex |
|---|---|
| **Sample preparation** | |
| Buffer | HEPES pH 7.5, NaCl, $CaCl_2$ |
| Concentrations (μM) | 0.65 fVa, 3.9 fXa, 1.3 prothrombin |
| Sample volume (μl) | 3 |
| Grid type | QuantiFoil (AU) R1/1 300 mesh |
| Glow discharge time (s) | 60 |
| Glow discharge current (mA) | 25 |
| Blotting chamber temp. (°C) | 4 |
| Blotting chamber humidity (%) | 100 |
| Blot time (s) | 1 |
| Blot force (N) | -7 |
| **Data collection and processing** | |
| Voltage (kV) | 300 |
| Detector mode | Counting – LZW-TIFF output |
| Indicated magnification | 130,000× |
| Pixel size (Å) | 0.829 |
| C2 aperture (μm) | 50 |
| Defocus range (μm) | -1.8 to -0.6 |
| Zero-loss slit width (eV) | No energy filter |
| Exposure rate (e/pix/s) | 12.63 |
| Exposure rate (e/Å$^2$/s) | 18.38 |
| Exposure time (s) | 2.72 |
| Total exposure (e-/Å$^2$) | 50.0 |
| Movies collected | 24,030 |
| **Refinement & validation** | |
| Symmetry imposed | C1 |
| Initial particle images (#) | 7,138,515 |
| Final particle images (#) | 76,039 |
| Map resolution (Å) | 3.11 |
| Model composition | |
| Non-hydrogen atoms | 18,403 |
| Protein atoms | 17,912 |
| Ions | 2 |
| Carbohydrate atoms | 489 |
| R.m.s. deviations | |
| Bond lengths (Å) | 0.004 |
| Bond angles (º) | 0.755 |
| Ramachandran favoured (%) | 89.92 |
| Ramachandran outliers (%) | 0.41 |
| MolProbity score | 2.36 |
| Clash score | 20.47 |
| Rotamer outliers (%) | 0 |

**Supplemental Table S2.** Cryo-EM data collection, refinement and validation statistics for the prothrombinase-meizothrombin complex (EMD- 56054; PDBID 9TLG).

| | Prothrombinase-Meizothrombin Complex |
|---|---|
| **Sample preparation** | |
| Buffer | HEPES pH 7.5, NaCl, $CaCl_2$ |
| Concentrations (µM) | 0.65 fVa, 3.9 fXa, 1.3 meizothrombin |
| Sample volume (µl) | 3 |
| Grid type | QuantiFoil (AU) R1/1 300 mesh |
| Glow discharge time (s) | 60 |
| Glow discharge current (mA) | 25 |
| Blotting chamber temp. (°C) | 4 |
| Blotting chamber humidity (%) | 100 |
| Blot time (s) | 1 |
| Blot force (N) | 0 |
| **Data collection and processing** | |
| Voltage (kV) | 300 |
| TEM mode | Counting – LZW-TIFF output |
| Indicated magnification | 165,000× |
| Pixel size (Å) | 0.729 |
| C2 aperture (µm) | 50 |
| Defocus range (µm) | -1.8 to -0.8 |
| Zero-loss slit width (eV) | 10 |
| Exposure rate (e/pix/s) | 6.09 |
| Exposure rate (e/Å$^2$/s) | 11.45 |
| Exposure time (s) | 4.39 |
| Total exposure (e-/Å$^2$) | 50.3 |
| Movies collected | 9,303 |
| **Refinement & validation** | |
| Symmetry imposed | C1 |
| Initial particle images (#) | 5,437,924 |
| Final particle images (#) | 25,376 |
| Map resolution (Å) | 3.08 |
| Model composition | |
| Non-hydrogen atoms | 16,552 |
| Protein atoms | 16,291 |
| Ions | 3 |
| Carbohydrate atoms | 258 |
| R.m.s. deviations | |
| Bond lengths (Å) | 0.003 |
| Bond angles (º) | 0.714 |
| Ramachandran favoured (%) | 89.39 |
| Ramachandran outliers (%) | 0.85 |
| MolProbity score | 3.02 |
| Clash score | 16.84 |
| Rotamer outliers (%) | 8.90 |

**Supplemental Table S3.** Data collection, refinement and validation statistics for PDBID 9T00. Highest resolution shell is in parenthesis.

| | Pre-2:a2-peptide |
|---|---|
| **Data collection** | |
| Space group | P1 |
| Cell dimensions | |
| *a*, *b*, *c* (Å) | 51.42, 51.57, 65.99 |
| α,β,γ (°) | 82.90, 85.33, 65.90 |
| Resolution (Å) | 38.76-2.80 (2.95-2.80) |
| $R_{merge}$ | 0.244 (1.03) |
| Mean $I/\sigma I$ | 3.0 (0.7) |
| $CC_{1/2}$ | 0.950 (0.423) |
| Completeness (%) | 99.2 (98.9) |
| Redundancy | 3.7 (3.4) |
| **Refinement & validation** | |
| Resolution (Å) | 38.76-2.80 |
| No. reflections | 14,261 |
| $R_{work}/ R_{free}$ | 0.259/0.305 |
| (2.87-2.80 Å) | (0.414/0.392) |
| No. atoms | |
| Protein/peptide | 5,114 |
| Water | 11 |
| B-factors | |
| All | 48.0 |
| R.m.s. deviations | |
| Bond lengths (Å) | 0.001 |
| Bond angles (º) | 0.785 |
| Ramachandran favoured (%) | 90.41 |
| Ramachandran outliers (%) | 1.46 |
| MolProbity score | 2.73 (82$^{nd}$ percentile) |

**Supplemental Table S4. Factor Xa (M17) light chain contacts with fVa.** Contacting residues identified by the program 'contact' are listed. Interaction type as identified by PISA are in the last column with HB for hydrogen bond and SB for salt-bridge.

| fXa | | fVa | | |
|---|---|---|---|---|
| Domain | Residue | Domain | Residue | Interaction Type |
| EGF2 | Met85 | A3 | Gln1629 | |
| | | | Thr1679 | |
| | | | Val1681 | |
| | Arg86 | A3 | Ser1652 | HB |
| | | | Ser1653 | |
| | | | Trp1665 | |
| | | | Glu1668 | SB |
| | | | Tyr1678 | HB |
| | | | Thr1679 | HB |
| | Leu88 | A3 | Glu1650 | |
| | | | Ser1652 | |
| | | | Phe1666 | |
| | | | Val1681 | |
| | | | His1683 | |
| | **Ser90Arg** | A2 | Asn629 | 2HB |
| | | A3 | Ser1648 | HB |
| | | | Tyr1649 | |
| | | | Glu1650 | |
| | | | His1683 | |
| | **Leu91Ala** | A3 | Val1627 | |
| | | | Arg1551 | |
| | | | His1683 | |
| | **Asp92Phe** | A3 | Asn1547 | |
| | | | Gly1549 | |
| | | | Asn1550 | |
| | | | Arg1551 | |
| | | | Val1627 | |
| | **Glu102Arg** | A3 | Glu1650 | 4SB |
| | | | Ser1652 | |
| | | | Phe1666 | |
| | **Glu103Val** | A3 | Trp1665 | |
| | Gln104 | A3 | Trp1665 | |
| | **Asn105Ser** | A3 | Trp1665 | |

**Supplemental Table S5. Factor Xa (M17) SP domain interactions with fVa**. Contacting residues identified by the program ‘contact’ are listed. Interaction type as identified by PISA are in the last column with HB for hydrogen bond and SB for salt-bridge. Chymotrypsin numbering used for SP, and M17 mutations are in bold.

| fXa | fVa | | |
|---|---|---|---|
| Chymo #ing | Domain | Residue | Interaction Type |
| Glu36 | a2-loop | Ser673 | |
| | | Thr674 | |
| | | Val675 | |
| Tyr60 | a2-loop | Phe668 | |
| Gln61 | a2-loop | Phe668 | |
| | | Glu669 | |
| | | Pro670 | |
| Ala61a | a2-loop | Ser673 | |
| Lys62 | a2-loop | Glu669 | |
| | | Pro670 | |
| | | Pro671 | HB |
| | | Glu672 | SB |
| | | Ser673 | |
| | | Thr674 | |
| Phe64 | a2-loop | Pro670 | |
| Lys65 | a2-loop | Thr674 | |
| Glu84 | a2-loop | Thr674 | |
| Val85 | a2-loop | Pro670 | |
| | | Pro671 | |
| Glu86 | a2-loop | Glu669 | |
| | | Pro670 | |
| | | Pro671 | |
| Val87 | a2-loop | Ile667 | |
| | | Phe668 | |
| | | Pro670 | |
| Val88 | a2-loop | Glu666 | |
| | | Ile667 | |
| | | Phe668 | 2HB |
| | | Glu669 | |
| | | Pro670 | |
| Ile89 | a2-loop | Tys665 | |
| | | Glu666 | |
| | | Ile667 | |
| Lys90 | a2-loop | Tys665 | |
| | | Glu666 | 2HB |
| | | Phe668 | |
| His91 | a2-loop | Ser664 | |
| | | Tys665 | |
| | | Glu666 | |
| Asn92 | a2-loop | Ser664 | |
| | | Tys665 | |
| | | Glu666 | |
| Lys109 | a2-loop | Pro671 | |
| Arg125 | A2 | Lys655 | |
| | | Cys656 | HB |

| | | | |
|---|---|---|---|
| | a2-loop | Ile657 | |
| | | Pro658 | |
| | | Asp659 | |
| Asp126 | A2 | Lys655 | |
| **Glu129Asn** | A2 | Phe576 | |
| | | Asp577 | |
| | | Lys655 | |
| **Ser130Glu** | A2 | Phe576 | |
| | | Val630 | |
| | | Lys655 | |
| Met131b | A2 | Asp577 | |
| **Thr132Lys** | A2 | Phe576 | |
| | | Asp577 | SB |
| | | Asp628 | 2SB |
| | A3 | Ser1546 | |
| Gln133 | A3 | Ser1546 | |
| Asp134 | A3 | Ser1546 | |
| Asp164 | A3 | Glu1686 | |
| Arg165 | A2 | Asp513 | 3SB |
| | | Asp577 | |
| | | Thr579 | |
| **Asn166His** | A2 | Gln509 | |
| | | Ala511 | HB |
| | | Ala512 | |
| | | Thr579 | |
| | | Gln581 | |
| | | Thr624 | |
| **Lys169Met** | A2 | Arg510 | HB |
| | | Ala511 | |
| | | Ala512 | |
| | | Asp513 | |
| | | Thr579 | |
| Leu170 | A2 | Gln471 | |
| | | Arg510 | HB |
| | | Ala511 | |
| Ser171 | A2 | Arg510 | |
| Ser172 | A2 | Arg510 | HB |
| **Ser173Asp** | A2 | Arg501 | 5SB |
| | | Arg510 | |
| Phe174 | A2 | Arg510 | HB |
| **Ile175Arg** | A2 | Asp504 | 3SB |
| | | Ile508 | |
| | | Gln509 | HB |
| | | Arg510 | |
| Ile176 | A2 | Arg510 | |
| | | Asp513 | |
| Gln178 | A2 | Asp513 | HB |
| | | Asp578 | |
| Thr232 | a2-loop | Asp659 | |
| **Ala233Arg** | a2-loop | Asp659 | 5SB |
| | | Glu662 | |
| Phe234 | a2-loop | Asp659 | |
| | | Glu662 | |

| | | | |
|---|---|---|---|
| Leu235 | a2-loop | Glu662 | |
| Lys236 | a2-loop | Pro658 | HB |
| | | Asp660 | HB |
| | | Glu662 | |
| Trp237 | a2-loop | Glu662 | |
| | | Asp663 | |
| | | Ser664 | |
| | | Tys665 | |
| Arg240 | a2-loop | Glu662 | |
| | | Asp663 | |
| | | Tys665 | HB |
| Ser241 | a2-loop | Tys665 | HB |
| Lys243 | a2-loop | Tys665 | |
| Thr244 | a2-loop | Tys665 | |
| | | Ile667 | |
| Gly246 | a2-loop | Ile667 | |

**Supplemental Table S6. Prothrombin interactions with fVa**. Contacting residues identified by the program 'contact' are listed. Interaction type as identified by PISA are in the last column with HB for hydrogen bond and SB for salt-bridge. Chymotrypsin numbering is given for residues in the SP domain.

| Prothrombin | | fVa | | |
|---|---|---|---|---|
| Mature #ing | Chymo #ing | Domain | Residue | Interaction Type |
| Glu253 | - | a1-loop | Arg313 | |
| Glu254 | - | a1-loop | Arg313 | |
| Glu255 | - | a1-loop | Arg313 | |
| Thr256 | - | a1-loop | Arg313 | |
| Gly257 | - | a1-loop | Arg313 | |
| Leu260 | - | a1-loop | Arg316 | |
| Asp261 | - | a1-loop | Arg316 | |
| | | a1-loop | Arg317 | |
| Arg266 | - | A2 | Arg321 | |
| Lys341 | 36 | a2-loop | Leu706 | |
| Gln344 | 38 | a2-loop | Leu706 | HB |
| | | | Gly707 | HB |
| | | | Ile708 | |
| | | | Arg709 | |
| Leu380 | 65 | a2-loop | Leu706 | |
| | | | Ile708 | |
| Arg382 | 67 | a2-loop | Ile708 | |
| | | | Arg709 | SB (C-term) |
| Lys385 | 70 | a2-loop | Arg709 | |
| Arg388 | | a2-loop | Arg709 | |
| Thr389 | 74 | a2-loop | Arg709 | |
| Arg390 | 75 | a2-loop | Arg709 | |
| Tyr391 | 76 | a2-loop | Gln699 | |
| | | | Asn700 | HB |
| | | | Ala703 | |
| | | | Ile708 | |
| | | | Arg709 | HB |
| Arg393 | 77a | a2-loop | His682 | 2HB |
| | | | Asp683 | |

| | | | | |
|---|---|---|---|---|
| | | | Arg684 | |
| | | | Leu685 | |
| | | | Tys696 | HB |
| | | | Asn700 | |
| Asn394 | 78 | a2-loop | Leu685 | |
| | | | Glu686 | |
| | | | Tys696 | |
| | | | Asn700 | |
| Glu396 | 80 | a2-loop | Gln699 | |
| Lys397 | 81 | a2-loop | Asp695 | |
| | | | Gln699 | |
| Ile398 | 82 | a2-loop | Tys698 | |
| | | | Gln699 | |
| | | | Leu702 | |
| | | | Ile708 | |
| Ser399 | 83 | a2-loop | Tys698 | |
| Met400 | 84 | a2-loop | Tys698 | |
| | | | Leu702 | |
| Lys427 | 110 | a2-loop | Asp695 | |
| | | | Tys698 | HB |
| Gln451 | 131 | a1-loop | Arg313 | HB |
| | | | Arg317 | |
| Ala452 | 132 | a1-loop | Arg317 | |
| Glu489 | 164 | a1-loop | Arg313 | |

**Supplemental Table S7. M17 SP domain interactions with prothrombin**. Contacting residues identified by the program 'contact' are listed. No HB or SB identified by PISA. Chymotrypsin numbering used for SP domain of fXa. Bold if not P4-P2′ (317-322). Note only 5 other residues contact fXa, so only 11 residues in total on prothrombin.

| fXa | Prothrombin | |
|---|---|---|
| Chymo #ing | Residue | Interaction Type |
| Phe41 | Ile321 | |
| | Val322 | |
| Cys42 | Arg320 | |
| | Ile321 | |
| His57 | Gly319 | |
| | Arg320 | |
| | Ile321 | |
| Cys58 | Ile321 | |
| Tyr60 | **Lys307** | |
| Lys96 | **Arg310** | |
| | Ile317 | |
| Glu97 | **Arg310** | |
| | Ile317 | |
| Tyr99 | Ile317 | |
| | Gly319 | |
| | Ile321 | |
| Arg143 | Val322 | |
| Lys148 | **Tyr510** | |
| Gln151 | Val322 | |
| Phe174 | **Ser315** | |

| | |
|---|---|
| | **Tyr316** |
| | Ile317 |
| Asp189 | Arg320 |
| Ala190 | Arg320 |
| Cys191 | Arg320 |
| Gln192 | Asp318 |
| | Gly319 |
| | Arg320 |
| | Ile321 |
| | Val322 |
| Gly193 | Arg320 |
| | Ile321 |
| | Val322 |
| Asp194 | Arg320 |
| Ala195 | Arg320 |
| | Ile321 |
| Val213 | Arg320 |
| Ser214 | Arg320 |
| Trp215 | Ile317 |
| | Asp318 |
| | Gly319 |
| | Arg320 |
| Gly216 | Asp318 |
| | Gly319 |
| | Arg320 |
| Glu217 | Asp318 |
| | Arg320 |
| Gly218 | Asp318 |
| | Arg320 |
| Arg222 | **Tyr316** |
| Tyr225 | Arg320 |
| Gly226 | Arg320 |
| Ile227 | Arg320 |
| Tyr228 | Arg320 |

**Supplemental Table S8. Prothrombin interactions with fXa**. Contacting residues identified by the program 'contact' are listed. No HB or SB identified by PISA. Chymotrypsin numbering used for SP domain of fXa.

| Prothrombin Residue | fXa Chymo #ing | Interaction Type |
|---|---|---|
| Lys307 | Tyr60 | |
| Arg310 | Lys96 | |
| | Glu97 | |
| Ser315 | Phe176 | |
| Tyr316 | Phe176 | |
| | Arg222 | |
| Ile317 | Lys96 | |
| | Glu97 | |
| | Tyr99 | |
| | Phe174 | |
| | Trp215 | |
| Asp318 | Gln192 | |

| | |
|---|---|
| | Trp215 |
| | Gly216 |
| | Glu217 |
| | Gly218 |
| Gly319 | His57 |
| | Tyr99 |
| | Gln192 |
| | Trp215 |
| | Gly216 |
| Arg320 | Cys42 |
| | His57 |
| | Asp189 |
| | Ala190 |
| | Cys191 |
| | Gln192 |
| | Gly193 |
| | Asp194 |
| | Ala195 |
| | Val213 |
| | Ser214 |
| | Trp215 |
| | Gly216 |
| | Glu217 |
| | Gly218 |
| | Tyr225 |
| | Gly226 |
| | Ile227 |
| | Tyr228 |
| Ile321 | Phe41 |
| | Cys42 |
| | His57 |
| | Cys58 |
| | Tyr99 |
| | Gln192 |
| | Gly193 |
| | Ala195 |
| Val322 | Phe41 |
| | Arg143 |
| | Gln151 |
| | Gln192 |
| | Gly193 |
| Tyr510 | Lys148 |

**Supplemental Table S9. Meizothrombin light chain (P) interactions with fVa**. Contacting residues identified by the program 'contact' are listed. 0 HB and 2 SB identified by PISA. Chymotrypsin numbering used for SP domain of fXa.

| Meizothrombin Residue | fVa Residue | Interaction Type |
|---|---|---|
| Asp261 | Glu314 | |
| Ser264 | Arg317 | |
| Asp265 | Arg317 | 2SB |
| Arg266 | His318 | |

**Supplemental Table S10. Meizothrombin light chain (P) interactions with fXa**.
Contacting residues identified by the program 'contact' are listed. 13 HB and 3SB identified by PISA. Chymotrypsin numbering used for SP domain of fXa.

| Meizothrombin | fXa | |
|---|---|---|
| Residue | Chymo #ing | Interaction Type |
| Arg266 | Asp173 | SB |
| | Phe174 | |
| Ala267 | Phe174 | |
| Ile268 | Lys96 | |
| | Glu97 | |
| | Thr98 | |
| | Tyr99 | |
| | Phe174 | |
| | Trp215 | |
| Glu269 | Tyr99 | |
| | Gln192 | HB |
| | Trp215 | |
| | Gly216 | HB |
| | Glu217 | |
| | Gly218 | HB |
| Gly270 | His57 | |
| | Tyr99 | |
| | Gln192 | HB |
| | Ser214 | |
| | Trp215 | |
| | Gly216 | |
| Arg271 | Phe41 | |
| | Cys42 | |
| | His57 | |
| | Asp189 | 3SB |
| | Ala190 | HB |
| | Cys191 | |
| | Gln192 | |
| | Gly193 | HB |
| | Asp194 | HB |
| | Ala195 | HB |
| | Val213 | |
| | Ser214 | |
| | Trp215 | |
| | Gly216 | |
| | Glu217 | |
| | Gly218 | HB |
| | Cys220 | |
| | Gly226 | |
| | Ile227 | |
| Thr272 | Phe41 | HB |
| | His42 | |
| | His57 | |
| | Cys58 | |
| | Gln61 | HB |

| | | |
|---|---|---|
| | Arg143 | |
| | Gln151 | |
| | Gln192 | |
| | Gly193 | |
| | Ala195 | |
| Ala273 | Phe41 | HB |
| | Gln61 | HB |
| | Gln151 | |
| | Gly193 | |
| Thr274 | Gln61 | |
| Ser275 | Gln61 | |
| Ser303 | Glu37 | |
| Arg310 | Arg150 | |
| Leu313 | Lys148 | |
| | Arg150 | |
| Glu314 | Arg150 | |
| Tyr316 | Lys148 | |
| Ile317 | His145 | |
| | Glu147 | |
| | Lys148 | |
| | Gly149 | |
| Asp318 | His145 | |
| Arg320 | Arg150 | |

**Supplemental Table S11. Meizothrombin SP domain (Q) interactions with fXa**. Contacting residues identified by the program 'contact' are listed. 3 HB and 0 SB identified by PISA. Chymotrypsin numbering used for SP domain of fXa.

| Meizothrombin | | fXa | |
|---|---|---|---|
| Residue | Chymo #ing | Chymo #ing | Interaction Type |
| 448 | Ser129b | Lys148 | HB |
| 449 | Leu129c | Lys148 | |
| 454 | Tyr134 | Lys148 | HB |
| 535 | Phe204a | Arg143 | HB |
| | | Lys148 | |
| | | Gly149 | |

**Supplemental Table S12.** F1 interactions with the SP in prothrombin. Contacting residues identified by the program 'contact' are listed. 8 HB and 5 SB identified by PISA. Chymotrypsin numbering used for SP domain of prothrombin.

| F1 | Prothrombin SP | | |
|---|---|---|---|
| Residue | Residue | Chymo #ing | Interaction Type |
| Glu85 | Arg498 | 173 | 2SB |
| Leu88 | Asp552 | 221 | |
| | Arg553 | 221a | |
| Arg90 | Trp468 | 147a | |
| | Thr469 | 147b | |
| | Asp552 | 221 | HB, 2SB |
| | Asp554 | 222 | SB |
| Ser91 | Trp468 | 147a | |

| | | | |
|---|---|---|---|
| | Trp547 | 215 | |
| | Asp552 | 221 | |
| Arg92 | Tyr367 | 60a | |
| | Trp370 | 60d | |
| | Glu466 | 146 | HB |
| | Thr467 | 147 | HB |
| | Trp468 | 147a | |
| | Thr469 | 147b | |
| | Ala470 | 147c | |
| Tyr93 | Tyr367 | 60a | |
| | Glu414 | 97a | |
| | Asn415 | 98 | |
| | Leu416 | 99 | HB |
| | Trp468 | 147a | |
| | Trp547 | 215 | |
| Pro94 | Tyr367 | 60a | HB |
| | Pro369 | 60c | |
| | Trp370 | 60d | |
| His95 | Tyr367 | 60a | |
| | Pro369 | 60c | |
| | Trp370 | 60d | |
| Lys96 | Pro369 | 60c | |
| | Trp370 | 60d | |
| Thr129 | Glu414 | 97a | |
| Asp130 | Arg413 | 97 | |
| | Glu414 | 97a | |
| Pro131 | Pro369 | 60c | |
| | Trp412 | 96 | |
| | Arg413 | 97 | |
| | Glu414 | 97a | |
| | Asn415 | 98 | |
| Thr132 | Trp412 | 96 | |
| | Arg413 | 97 | 2HB |
| Arg134 | Pro369 | 60c | HB |
| | Trp370 | 60d | |

**Supplemental Table S13.** F1 clashes (<2.3Å) with the SP in meizothrombin. Contacting residues identified by the program 'contact' are listed. 0 HB and 2 SB identified by PISA. Chymotrypsin numbering used for SP domain of meizothrombin.

| F1 | Meizothrombin SP | | |
|---|---|---|---|
| Residue | Residue | Chymo #ing | distance (Å) |
| Glu85 | Arg498 | 173 | 1.57 |
| Arg92 | Trp370 | 60d | 1.31 |
| Tyr93 | Leu416 | 99 | 1.70 |
| His95 | Trp370 | 60d | 1.52 |
| Lys96 | Trp370 | 60d | 1.26 |
| Pro131 | Arg413 | 97 | 1.58 |
| Arg134 | Pro369 | 60c | 1.83 |

**Supplemental Table S14.** K2 contacts (<5Å) with the SP in prothrombin. Contacting residues identified by the program 'contact' are listed. 1,879.8 $Å^2$ buried; 3 HB and 15 SB identified by PISA. Chymotrypsin numbering used for SP domain.

| K2 | Prothrombin SP | | |
|---|---|---|---|
| Residue | Residue | Chymo #ing | Interaction Type |
| Arg174 | Arg418 | 101 | |
| | Asp503 | 178 | 4SB |
| | Arg565 | 233 | |
| Thr186 | Pro408 | 92 | |
| | Arg409 | 93 | |
| | Asn411 | 95 | |
| His187 | Pro408 | 92 | |
| | Tyr410 | 94 | |
| | Asn411 | 95 | |
| | Arg413 | 97 | |
| His205 | Gln571 | 239 | |
| | Asp575 | 243 | SB |
| Gln206 | Gln571 | 239 | |
| | Asp575 | 243 | |
| Asp223 | Lys568 | 236 | |
| Gly224 | Lys568 | 236 | |
| Asp225 | Arg565 | 233 | |
| | Lys568 | 236 | |
| | Trp569 | 237 | |
| | Lys572 | 240 | SB |
| Glu226 | Arg443 | 126 | |
| | Phe564 | 232 | |
| | Arg565 | 233 | SB |
| | Lys568 | 236 | |
| Glu227 | His407 | 91 | |
| | Arg409 | 93 | |
| | Arg418 | 101 | 4SB |
| | Asn504 | 179 | HB |
| | Arg565 | 233 | SB |
| Trp230 | Lys572 | 240 | |
| Lys236 | Glu579 | 247 | |
| Pro237 | Gln576 | 244 | |
| | Glu579 | 247 | |
| Gly238 | Gln576 | 244 | |
| Asp239 | Gln576 | 244 | |
| Phe240 | Pro408 | 92 | |
| | Trp569 | 237 | |
| | Lys572 | 240 | |
| | Val573 | 241 | |
| Gly241 | Pro408 | 92 | |
| Tyr242 | His407 | 91 | HB |
| | Pro408 | 92 | |
| | Arg409 | 93 | |
| | Trp569 | 237 | |
| Asp244 | Arg409 | 93 | SB |
| Cys248 | Asp503 | 178 | |
| Glu249 | Arg490 | 165 | SB |
| | Lys494 | 169 | SB |

| | | | |
|---|---|---|---|
| | Ile501 | 176 | |
| | Thr502 | 177 | |
| | Asp503 | 178 | |
| Glu250 | Asp503 | 178 | HB |
| | Arg565 | 233 | |

**Supplemental Table S15.** K2 contacts (<5Å) with the SP in meizothrombin. Contacting residues identified by the program 'contact' are listed. 1,579.1 Å$^2$ buried; 8 HB and 10 SB identified by PISA. Chymotrypsin numbering used for SP domain.

| K2 | Meizothrombin SP | | |
|---|---|---|---|
| Residue | Residue | Chymo #ing | Interaction Type |
| Leu189 | Arg413 | 97 | |
| Leu202 | Pro408 | 92 | |
| Lys204 | Lys572 | 240 | HB |
| | Gln576 | 244 | |
| His205 | Trp569 | 237 | |
| | Lys572 | 240 | HB |
| | Gln576 | 244 | |
| Gln206 | His407 | 91 | |
| | Pro408 | 92 | |
| | Arg409 | 93 | |
| | Trp569 | 237 | |
| | Lys572 | 240 | HB |
| Asp207 | Lys568 | 236 | |
| Asp223 | Arg409 | 93 | |
| | Arg418 | 101 | |
| Asp225 | Arg409 | 93 | SB |
| | Asp417 | 100 | |
| | Arg418 | 101 | 3SB |
| | Thr502 | 177 | |
| Glu226 | Arg490 | 165 | |
| | Arg500 | 175 | |
| | Ile501 | 176 | |
| | Thr502 | 177 | |
| | Asp503 | 178 | HB |
| Glu227 | Glu414 | 97a | |
| | Arg500 | 175 | 4SB |
| | Ile501 | 176 | |
| | Thr502 | 177 | |
| Trp230 | Arg409 | 93 | |
| Tyr232 | Pro408 | 92 | |
| | Arg409 | 93 | |
| Pro237 | Leu366 | 60 | |
| | Ile406 | 90 | |
| | His407 | 91 | |
| | Pro408 | 92 | |
| | Tyr410 | 94 | |
| | Trp412 | 96 | |
| Gly238 | His407 | 91 | |
| | Pro408 | 92 | HB |
| | Tyr410 | 94 | |
| | Asn411 | 95 | |

| | | | |
|---|---|---|---|
| | Trp412 | 96 | |
| | Arg413 | 97 | HB |
| Asp239 | Pro408 | 92 | |
| | Trp412 | 96 | |
| | Arg413 | 97 | 2SB |
| Phe240 | Pro408 | 92 | |
| | Arg409 | 93 | |
| | Tyr410 | 94 | |
| | Asn411 | 95 | |
| | Arg413 | 97 | |
| | Arg418 | 101 | |
| Tyr242 | Arg409 | 93 | |
| | Asn411 | 95 | HB |
| | Glu414 | 97a | HB |
| | Asp417 | 100 | |
| | Arg500 | 175 | |

**Supplemental Table S16.** K2 in prothrombin position clashes (<2.3Å) with the SP in meizothrombin. Contacting residues identified by the program 'contact' are listed. Chymotrypsin numbering used for SP domain of meizothrombin.

| K2 | Meizothrombin SP | | |
|---|---|---|---|
| Residue | Residue | Chymo #ing | distance (Å) |
| Thr186 | Arg409 | 93 | 1.37 |
| Ser203 | Gln576 | 244 | 2.23 |
| Asp225 | Lys568 | 236 | 2.17 |
| Phe240 | Trp569 | 237 | 2.20 |
| Tyr242 | His407 | 91 | 1.29 |
| Glu249 | Arg490 | 165 | 1.98 |
| Glu250 | Arg565 | 233 | 2.26 |
